# N-terminal Trimming Supports Proteostasis Under Secretory Protein Mislocalization Stress

**DOI:** 10.64898/2026.10.13.756067

**Authors:** Jasmine P Castellanos, Khanh K. Nguyen, Qiqi Lin, Ziqi Lyu, Zi Yang, Fabella Sarwono, Jean Carlo Gonzalez, Walter Nguyen, Lyna T. Huynh, Carolina Canizal, Prasanna Satpute-Krishnan, Wei Shi, Joseph C. Genereux

## Abstract

Secretory proteins that fail to enter the endoplasmic reticulum (ER) are mislocalized to the cytosol, where they threaten cellular proteostasis. How cells manage these mislocalized proteins remains poorly understood. Here, we show that inefficient Sec61 translocon activity induces a transcriptional stress response, which is enhanced by blocking NRF-1-dependent adaptation of proteasomal capacity. To maintain proteasomal capacity, thermodynamically stable mislocalized proteins persist in the cytosol of HepG2 and HEK293T cells, while mislocalized proteins that are inherently more aggregation-prone are preferentially ubiquitinated and degraded. This triaging process is accomplished through N-terminal trimming of the signal sequence of thermodynamically stable proteins, which simultaneously reduces their aggregation and prevents proteasomal degradation. These findings reveal a stability-dependent cytosolic triaging mechanism that matches substrate burden to proteasome capacity when ER import is challenged.

**TOC Graphic:** 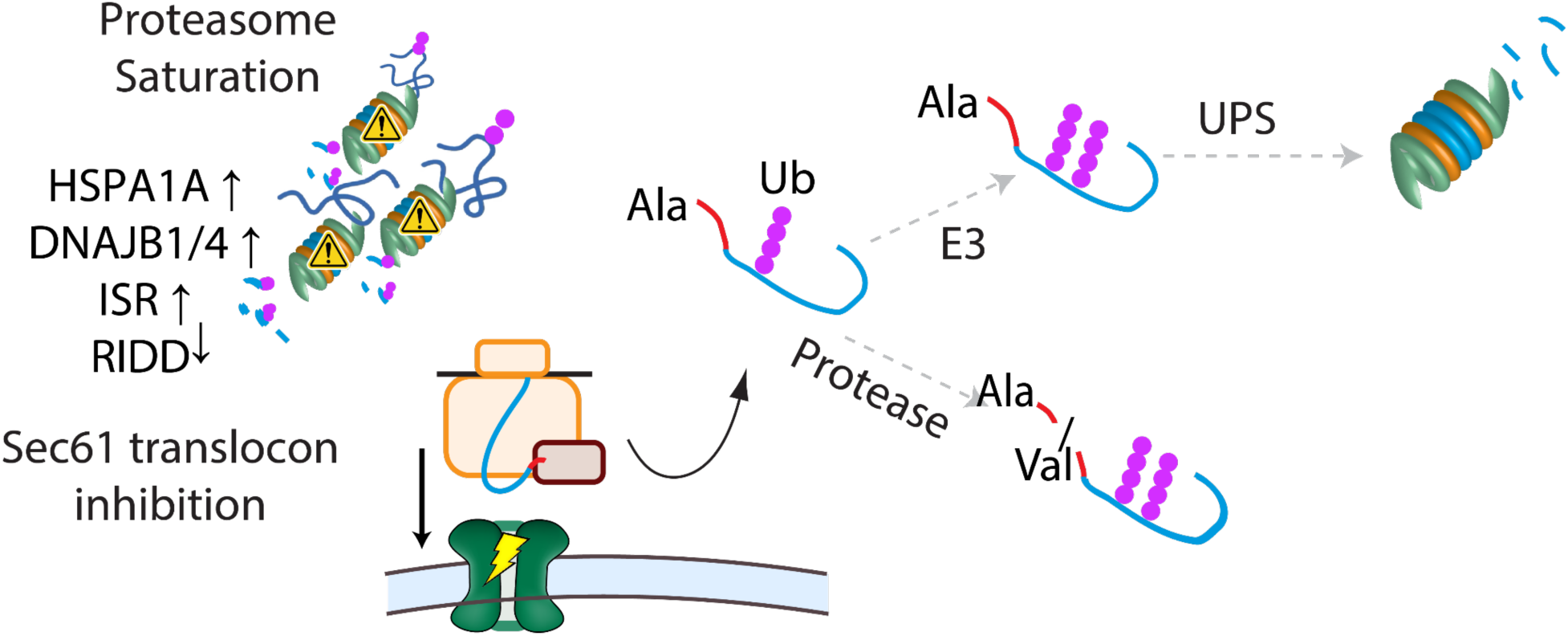

## Background

Correct intracellular localization of proteins plays an essential role in human health.^1–3^ A study of human missense variants across 1,269 disorder-related proteins revealed that mislocalization of secretory pathway proteins is a common characteristic of pathogenic variants.^2^ Most secretory pathway substrates, including secreted or membrane proteins, are co-translationally translocated into to the ER through a signal sequence and then fold in the ER.^1,2,4,5^ There are multiple ways that this targeting can fail. For example, in insulin-secreting pancreatic β cells, signal sequence mutations in proinsulin impair efficient translocation into the ER, leading to aggregation in the cytosol, cytosolic stress, and cell death. These cellular events are ultimately associated with type 2 diabetes phenotypes.^1^ A variety of environmental toxicants and emerging therapeutic molecules inhibit ER translocation through direct binding and inhibition of the Sec61 translocon.^6–10^ ER stress may also dysregulate protein translocation into the ER through pre-emptive quality control (ER preQC), leading to proteins becoming ‘orphaned’ in the cytosol. In the case of ER preQC, mislocalization has been proposed as a protective mechanism to reduce the protein folding burden in the ER.^11^ However, cellular prion (PrP^C^) misprocessing such as observed during ER preQC leads to mislocalization and subsequent degradation, and has been linked to neurodegenerative disease.^3^ Additionally, we recently found that disrupting the Sec61-associated protein Hspa13 leads to accumulation of transthyretin (TTR; an established preQC substrate) in the cytosol.^12^

ER stress induces the unfolded protein response (UPR) though the ER transmembrane receptor proteins PERK, IRE1, and ATF6.^13,14^ One study found that IRE-1 dependent Derlin upregulation is responsible for decreased Sec61 translocon activity during ER preQC.^13^ In contrast, another study revealed that phosphorylation of eIF2α by PERK downregulates signal recognition particle 14 (SRP14), thus decreasing targeting of newly synthesized ER proteins to Sec61 translocon.^14^ During ERpreQC mislocalized PrP^C^ is recognized by molecular chaperone Bag6, ubiquitinated by the E3 ligase RNF126, and then degraded by the ubiquitin-proteasome system (UPS).^15,16^ However, Bag6 recognition may not be universal across mislocalized proteins, given that Bag6 primarily recognizes membrane proteins^3,16,17^ and is sensitive to specific hydrophobic regions on its substrates.^18^ Hence, it is currently unclear what factors guide the fate of mislocalized secreted proteins.

Herein, we use Sec61 translocon inhibitors and ER stressors to induce protein mislocalization of canonical secretory pathway substrates in human tissue culture, and investigate how this burden impacts cellular proteostasis. We find that inhibition of ER import induces acute proteostasis stress and challenges proteasomal capacity. Further, while highly destabilized mislocalized proteins are rapidly degraded through UPS, less destabilized proteins evade degradation and persist in the cytosol. We show that these persistent substrates undergo N-terminal trimming, which reduces hydrophobicity, limits aggregation, and alleviates proteasomal overload during impaired ER import.

## Results

### Mislocalized Secretory Proteins Challenge Proteostasis and Proteasomal Capacity

Sec61 translocon inhibition induces acute mislocalization of secretory proteins to the cytosol. The mislocalized proteins will either be degraded, taxing proteasomal capacity, or accumulate, potentially aggregating and challenging cytosolic proteostasis. At the same time, ER proteostasis could either be enhanced due to a decrease in nascent unfolded proteins or challenged through remodeling of Sec61 membrane complexes.^19–21^ To induce mislocalization, we used two structurally distinct Sec61 inhibitors: the widely employed mycolactone (mycA/B)^22–24^ and the ipomoeassin F analog SW3,^10,25,26^ which engage the Sec61 translocon through different contacts at the lateral gate.^7^ Both compounds inhibit ER import of TTR with similar potency (SW3 EC₅₀ ≈ 2 nM; **Supplemental Figure S1A**) supporting their use as complementary tools to disrupt Sec61 translocation. MycA/B is highly unstable, adheres to plastics, and requires a multi-step synthesis.^27,28^ SW3 efficiently inhibits Sec61 with greater stability and a simpler synthesis than mycA/B.

To assess cellular stress during inadequate Sec61 translocon capacity, we inhibited Sec61 in hepatocyte cells, which manage a high secretory load, and measured poly-A tailed RNA transcripts. When proteasomal capacity becomes limited, homeostasis is restored through production of active NRF-1 by the deglycosylase NGLY1. We included conditions with the NGLY1 inhibitor WRR139 to prevent this enhancement of proteasomal capacity during stress (**Figure 1A**). Sec61 inhibition by either mycA/B or SW3 leads to a moderate transcriptional profile, with 433 differentially expressed genes following mycA/B treatment and 91 differentially expressed genes following SW3 treatment (q < 0.05, |log_2_FC| ≥ 1) (**Supplemental Figure S1B, S1C**). The transcriptional signatures associated with each treatment are comparable (Figure 1B); for example, major downregulated pathways include those associated with mitosis and cellular growth, indicating general toxicity (**Supplemental Figure S1D, S1E**). SLC3A2 is a subunit of the CD98 amino acid transporter and its abundance is known to increase following mycA/B treatment.^23,29^ Consistent with those studies, *SLC3A2* increases 6.43 ± 0.57-fold with SW3 treatment and 9.12 ± 0.42-fold with mycA/B. To understand how stress pathways were impacted by Sec61 inhibition, we focused on gene signatures that serve as the most selective for the major cellular stress pathways relevant to proteostasis.^30^ The effects of SW3 and mycA/B on stress responses are nearly identical (**Figure 1B**). Previous studies had found that Sec61 translocon inhibition increases *HSPA1A* levels, suggesting heat shock response (HSR) activation.^22,23^ While we see significant induction of canonical HSR targets by both Sec61 inhibitors, the induction is moderate (**Figure 1C**) compared to the well-characterized HSR induction by proteasome inhibitors, such as bortezomib (Btz, **Supplemental Figure S1F**).^31,32^ At the protein level, HSR target chaperones HSP70 and DNAJB1 are strongly induced by SW3, with a stronger induction when SW3 is combined with WRR139 (**Figure 1D**), showing that ER import inhibition remodels the cytosolic chaperone environment through decreased proteasomal capacity.

**Figure 1.**
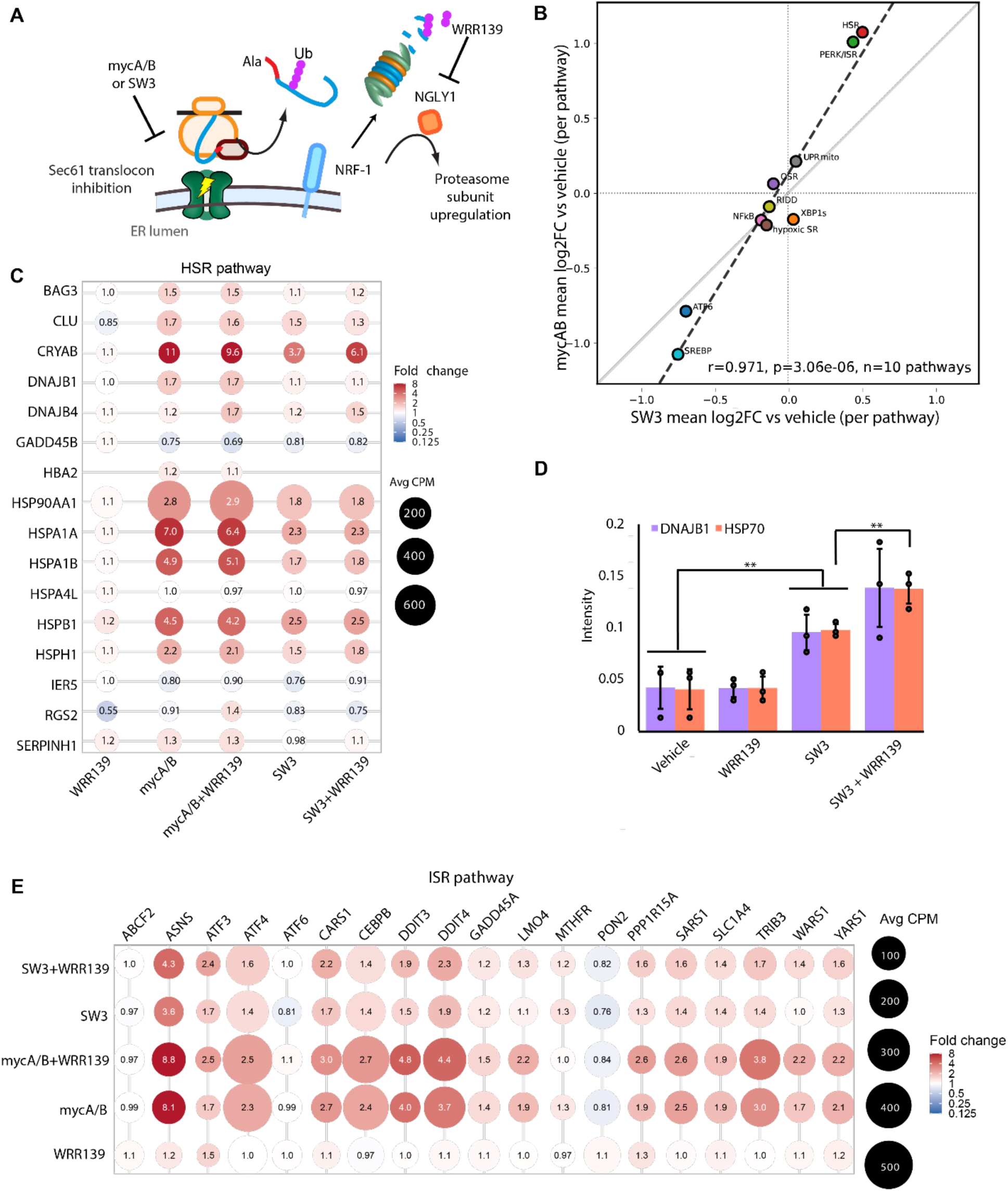
Mislocalized secretory protein increases cellular stress by overwhelming proteasome capacity. **A.** Schematic of RNAseq experiment and conditions. **B.** Pathway-averaged fold-change (log2FC) correlations comparing SW3 to mycAB across 10 stress-response pathways. Each point represents the mean log2FC of all marker genes within that pathway (n=10 pathways). **C.** Fold change in mRNA expression relative to vehicle control for target genes as determined by RNAseq for HSR target genes in HepG2 cells treated for 16 h with 50 nM SW3, 25 nM mycA/B, 10 µM WRR139, or vehicle as indicated. Each dot represents one gene under one condition. Color and printed value indicates fold change relative to vehicle (blue = decreased, red = increased). Dot size reflects mean cpm for that gene under that condition (log10 scale), as a measure of relative expression level/read depth. log2FC was calculated as log2(mean of 3 treatment replicates / mean of 3 vehicle replicates), computed independently within each dataset. For visualization, (log2FC values were converted to linear fold change (FC = 2^log2FC^) for the values displayed on each bubble and the colorbar scale; dot color remains mapped to log2FC to preserve a symmetric, equal-weighted representation of up- and down-regulation. **D.** Quantification of SDS-PAGE-separated lysates from cells treated for 16 h with 10 nM SW3, 10 µM WRR139, or vehicle as indicated. Error bars represent S.D. **E.** Fold change back-calculated from log2FC in mRNA expression relative to vehicle control as determined by RNAseq for ISR targets genes in HepG2 cells treated for 16 h with 50 nM SW3, 25 nM mycA/B, 10 µM WRR139, or vehicle as indicated.

IRE1 activation during the UPR induces both RIDD and a transcriptional response mediated by activation of the transcription factor XBP1s. Most reports have found that XBP1s and RIDD are activated during Sec61 inhibition, with one report finding XBP1s is not activated by mycA/B.^10,22,29,33,34^ We do not see XBP1s activation by SW3 or mycA/B, though there is mild reduction of RIDD targets (**Supplemental Figure S1G**).^35^ By contrast, Sec61 inhibition alongside WRR139 treatment upregulates a broad set of XBP1s target genes, consistent with canonical IRE1 activation (**Supplemental Figure S1G-H**). Neither SW3 nor mycA/B activate ATF6 genes, indicating that the UPR is not generally activated (**Supplemental Figure S1I**). Rather, a broad suppression of ATF6 target genes was observed, mirroring our findings with another S1P/S2P substrate, SREBP, (**Figure 1B and Supplemental Figure S1I**) and suggesting a common mechanism of restricted protein flux into the ER.^10,36^ WRR139 consistently attenuates ATF6 pathway downregulation across nearly every gene in the panel under both Sec61 inhibitors (**Supplemental Figure S1I**); though the pathway remains net negative overall, with only a few genes (e.g. DNAJB11, HYOU1, MIS12) crossing into net-positive territory (**Supplemental Figure S1I**). SW3 mildly activates the ISR (44.3% increase), while mycA/B activates it more strongly (138.6% increase), as indicated by induction of target genes such as ASNS and DDIT4 (**Figure 1E**). In summary, the transcriptional profile of Sec61 inhibition is consistent with stress in both the ER and the cytosol.

Strikingly, despite WRR139 having little effect on the transcriptome on its own, the combination of WRR139 with SW3 or mycA/B induces stronger responses for all of the stressors described herein (**Figures 1C, 1E, and Supplemental Figure S1H**). These effects indicate that cellular stress by impaired ER import is largely driven by proteasomal capacity being challenged by mislocalized protein. In the case of CHOP/DDIT3, a transcription factor responsible for regulating apoptosis, SW3 alone increases expression (FC = 1.48 ± 0.07 S.E.M), and SW3+WRR139 co-treatment increases it further (FC = 1.92 ± 0.17). DDIT3 expression under mycA/B treatment is strongly elevated (FC = 4.03 ± 0.08), with mycA/B+WRR139 co-treatment producing an additional increase (FC = 4.76 ± 0.30). By contrast, WRR139 has far less effect on the transcriptional program associated with Btz treatment (**Supplemental Figure S1J-K**); presumably the ability to enhance proteasomal capacity has little consequence when any newly synthesized proteasomes are inhibited. This suggests that while the NRF1 stress response is partially sufficient to maintain proteostasis during Sec61 inhibition, in its absence the proteasome machinery is overwhelmed.

### Mislocalized Secretory Proteins Persist in the Cytosol

Having found that proteasomal capacity becomes limiting during Sec61 inhibition, we examined the fate of individual mislocalized secretory proteins. We focused on two established ER preQC substrate proteins that are commonly used to study ER protein import, folding, and secretion. Both the wild-type protein and nearly 200 variants of TTR are associated with amyloidosis, ^19,20,37,38^ while folding-incompetent and aggregation-prone variants of alpha-1-antitrypsin (A1AT) lead to loss-of-function and gain-of-function phenotypes.^39^ Neither TTR nor A1AT is endogenously expressed in HEK293T cells.^37,41,42^ Therefore, to investigate their fate under conditions of impaired ER import, we transfected HEK293T cells with either TTR or A1AT expression plasmids. In immunoblots of lysates from HEK293T cells overexpressing FLAG-tagged TTR (^FLAG^TTR), a single ∼17-kDa band is observed, consistent with efficient ER entry and signal sequence cleavage, hereafter referred to as the mature form (**Figure 2A lane 1 and Figure 2B**). After treatment with 25 nM mycA/B for 16 h, the mature form is no longer detected. However, two immature ^FLAG^TTR isoforms are observed as distinct slow- and fast-migrating bands, both migrating more slowly than the mature form (**Figure 2A, lane 2**). These slow- and fast-migrating bands are similar to the two immature TTR forms that we observed previously upon *HSPA13* overexpression.^12^ The immature slow- and fast-migrating isoforms respond differently to proteasomal inhibition by 1 µM MG132 (**Figure 2A, lane 3**). The slow-migrating isoform is unaffected, while the fast-migrating isoform is slightly protected by proteasomal inhibition. Similar results were obtained with the alternative proteasome inhibitor Btz (**Supplemental Figure S2A**). The persistence of mislocalized TTR contrasts with the previously reported model of mislocalized secretory protein clearance, wherein mislocalized proteins are rapidly ubiquitinated and proteasomally degraded.^13,15,16,43^ Notably, HepG2 liver cells, which endogenously express and secrete TTR,^44^ upon mycA/B treatment also accumulate two slow- and fast-migrating immature TTR isoforms (**Figure 2C and Supplemental Figure S2B**), with the fast-migrating form protected by proteasome inhibition. Consistent with Sec61 translocon inhibition, SW3 treatment also produced the persistent slow-migrating ^FLAG^TTR isoform and the proteasomally degraded fast-migrating ^FLAG^TTR isoform in a similar manner as mycA/B (**Supplemental Figure S2C lane 5**).

**Figure 2A:**
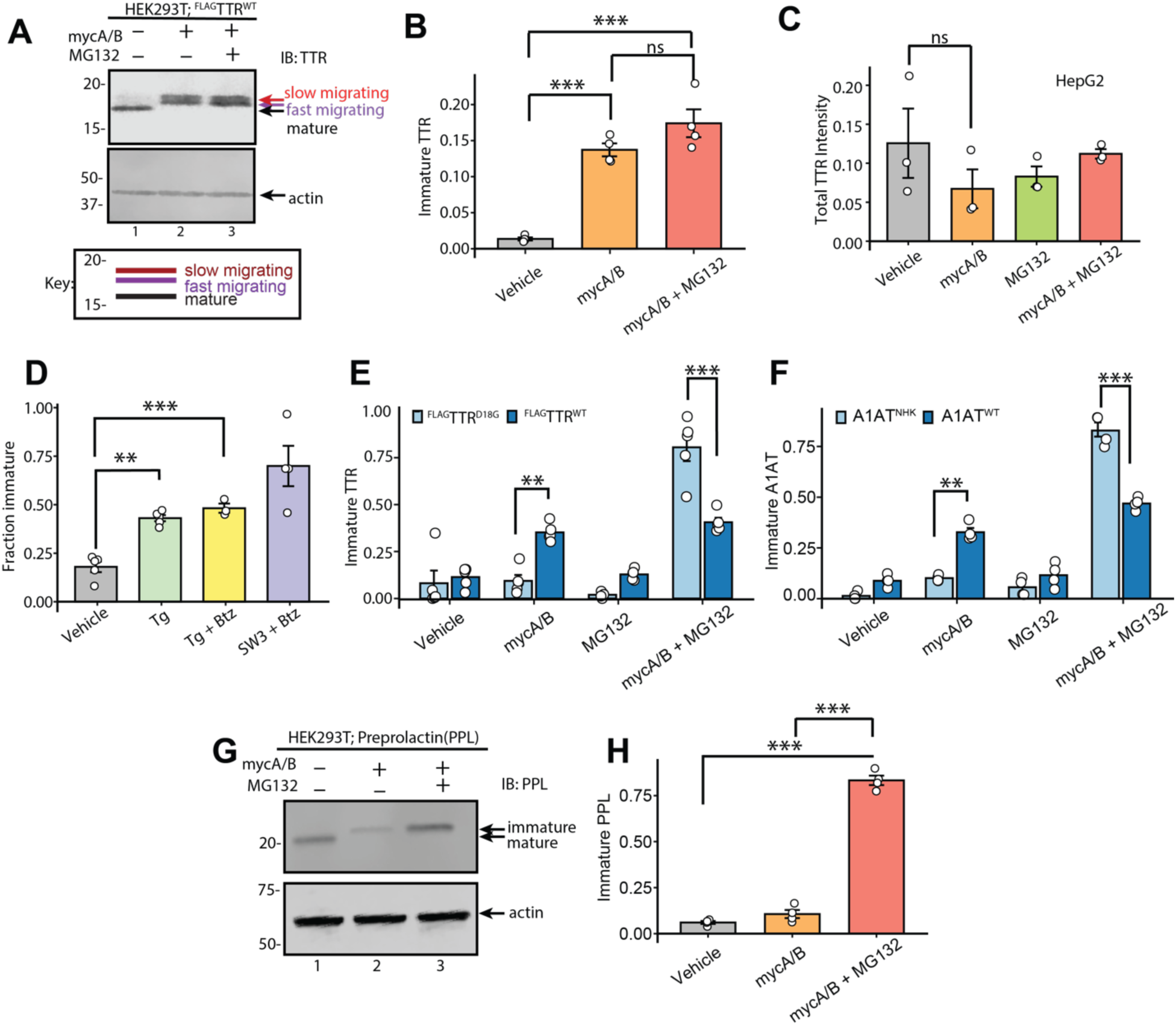
Mislocalized TTR produces a persistent immature isoform. Representative immunoblot of SDS-PAGE-separated lysates from HEK239T cells overexpressing ^FLAG^TTR and treated as indicated (25 nM mycA/B 16 h; 1 µM MG132 16 h). Actin is also blotted as loading control. Key for the migration of isoforms is below the blot. **B.** Quantification of immunoblots of HEK293T cells expressing ^FLAG^TTR^WT^ lysates under the indicated conditions (Repeated-measures ANOVA, F = 68.93 > Fcrit = 5.14, p < 0.001; n = 4 independent experiments. Post hoc comparisons (Tukey-adjusted). Densitometric values were normalized to the sum of the whole blot intensity. **C.** Quantification of endogenously expressed TTR in HepG2 treated as indicated (25 nM mycA/B 16 h; 1 µM MG132 16 h; *n* = 3). Actin also blotted as loading control. Densitometric values were normalized to the average intensity of TTR in control. **D.** Quantification immunoblots of SDS-PAGE-separated lysates from HEK239T cells overexpressing ^FLAG^TTR (50 nM Tg 16 h, *n* = 4). A significant outlier was identified through Grubbs test (α = 0.05) in the Tg+Btz group (*G* = 1.469, *p* = 0.0406), with the most extreme value being 0.0809 and were removed before performing statistical comparisons. Tg suppresses protein synthesis; the immature fraction is normalized to total protein. **E.** Quantification of immunoblot of SDS-PAGE-separated lysates from HEK239T cells overexpressing ^FLAG^TTR^WT^ or ^FLAG^TTR^D18G^ and treated for 16 h as indicated (25 nM mycA/B; 1 µM MG132; 100 µM chloroquine) (ANOVA *F* = 43.41 < *F*_crit_ = 2.31; *n* = 5). Densitometric values were normalized to the sum of the whole blot intensity. **F.** Quantification of immunoblot of SDS-PAGE-separated lysates from HEK239T cells overexpressing ^FLAG^A1AT^WT^ or ^FLAG^A1AT^NHK^ and treated for 16 h as indicated (25 nM mycA/B; 1 µM MG132) (ANOVA *F* = 171.3 < F_crit_ = 2.42; *n* = 4). ^FLAG^A1AT^NHK^ migrates faster than ^FLAG^A1AT^WT^ due to a frameshift truncation. **G.** Representative immunoblot of SDS-PAGE-separated lysates from HEK239T cells overexpressing preprolactin treated as indicated (25 nM mycA/B 16 h; 1 µM MG132 16 h). **H.** Quantification of immunoblot G. Densitometric values were normalized to the sum of the whole blot intensity. Significance codes: *** p < 0.001; ** p < 0.01; * p < 0.05. Unless otherwise noted, data are presented as mean ± SD.

ER preQC is induced by ER stressors, such as the SERCA inhibitor thapsigargin (Tg). Tg treatment induces the appearance of the slow-migrating immature isoform (**Supplemental Figure S2C, lane 2**), consistent with its reported ability to induce preQC^13,45^. As seen with Sec61 translocon inhibition, the combination of Tg treatment and proteasomal inhibition yields both the slow- and fast-migrating ^FLAG^TTR isoforms (**Supplemental Figure S2C, lane 3**). However, the mature ER-processed ^FLAG^TTR is still present (**Supplemental Figure S2C, lane 3 versus 5**), consistent with Tg producing only partial attenuation of ER import (**Figure 2D**). These results show that lower Sec61 translocon activity generally leads to the persistence of two distinct immature TTR isoforms, and this mislocalized TTR is mostly resistant to degradation.

Protein quality control mechanisms canonically recognize misfolded proteins and redirect them towards modification, segregation, and/or degradation.^46,47^ We considered that a strongly amyloidogenic TTR variant might be handled differently upon mislocalization. The D18G variant of TTR is highly destabilized and undergoes efficient aggregation that leads to amyloidosis in patients.^48^ We compared the degradation of mislocalized ^FLAG^TTR^D18G^ to ^FLAG^TTR^WT^ in HEK239T cells by immunoblot (**Figure 2E and Supplemental Figure S2D**). ^FLAG^TTR^D18G^ band intensity decreases in cells treated with mycA/B alone (**Supplemental Figure S2D, lane 3**), but band intensity returns upon proteasomal inhibition with MG132 (**Supplemental Figure S2D, lane 4**). Treatment with mycA/B and MG132 results in accumulation of both the fast- and slow-migrating isoforms of ^FLAG^TTR^WT^(**Supplemental Figure S2D, lane 8**), while only the fast-migrating isoform is produced for ^FLAG^TTR^D18G^ (**Supplemental Figure S2D, lane 4**). These results indicate that mislocalized TTR is subject to quality control, where stable variants accumulate and unstable variants are degraded.

To evaluate the generality of this finding, we compared A1AT^WT^ to its highly destabilized variant Null Hong Kong (A1AT^NHK^).^40,49^ A1AT has three N-linked glycan sites, and hence inhibiting its ER import should lower the apparent molecular weight by preventing glycosylation of A1AT in the ER. The migration of non-glycosylated A1AT was validated through PNGase treatment (**Supplemental Figure S2E**). A1AT^WT^ is persistent even when mislocalized to the cytosol by mycA/B treatment, while A1AT^NHK^ is degraded upon mislocalization (**Figure 2F and Supplemental Figure S2F, lane 3 and lane 7**). Proteasome inhibition significantly restores the immature A1AT^NHK^, indicating that this degradation is through UPS (**Supplemental Figure S2F, lane 8**). Thus, for both TTR and A1AT upon mislocalization, stable proteins can persist in the cytosol, but destabilized variants are efficiently targeted for degradation.

The ER contains protein disulfide isomerases (PDIs), which catalyze disulfide bond formation.^50^ Sec61 translocon inhibition prevents secretory proteins from entering the ER and therefore accessing these oxidative folding factors. To test whether dependence on ER-localized oxidative folding influences the fate of mislocalized secretory proteins, we examined secretory protein preprolactin (PPL), which requires ER-localized oxidative folding and directly interacts with PDI.^51–53^ PPL lacks the capacity to fold in the reducing cytosolic environment, we predicted that PPL would be intrinsically unstable when mislocalized. Consistent with this prediction, mislocalized PPL is less abundant under Sec61 inhibition, and proteasome inhibition rescues PPL abundance (**Figure 2G, H**). These findings support a shared mechanism where quality control triages mislocalized secretory proteins on the basis of their folding stability.

### Cytosolic N-terminal Trimming Protects Mislocalized TTR^WT^ from Degradation

Quality control pathways typically recognize misfolded proteins for modification (e.g. by ubiquitin or mannose trimming) and subsequent degradation. We used mass spectrometry to identify the modifications driving differential handling of the two immature TTR isoforms. We isolated samples enriched in the distinct TTR isoforms using the following conditions: ^FLAG^TTR^D18G^ alone to yield the mature protein, ^FLAG^TTR^D18G^ with mycA/B and MG132 to yield the fast-migrating isoform, and ^FLAG^TTR^WT^ with mycA/B alone to generate the slow-migrating isoform. These isoforms were immunopurified, yielding clean preparations by silver stain analysis of SDS-PAGE-purified eluates (**Figure 3A**), and analyzed by mass spectrometry. Mature ^FLAG^TTR^D18G^ is observed at the expected mass for signal sequence cleavage, 15,693 Da. The fast-migrating band has a mass of ∼17,687 Da, corresponding to the full-length cytosolic precursor ^FLAG^TTR^D18G^ with the signal sequence intact and methionine (Met) removal. A minor peak at ∼17,818 Da is consistent with initiator methionine retained, and a third precursor at ∼17,729 Da reflects Met removal and N-terminal acetylation (see **Supplemental Figure S3A1-3)**. Together, these peaks represent normal N-terminal processing of a signal-bearing precursor that has been excluded from ER import. We will now refer to this group collectively as the intact isoform.

**Figure 3A.**
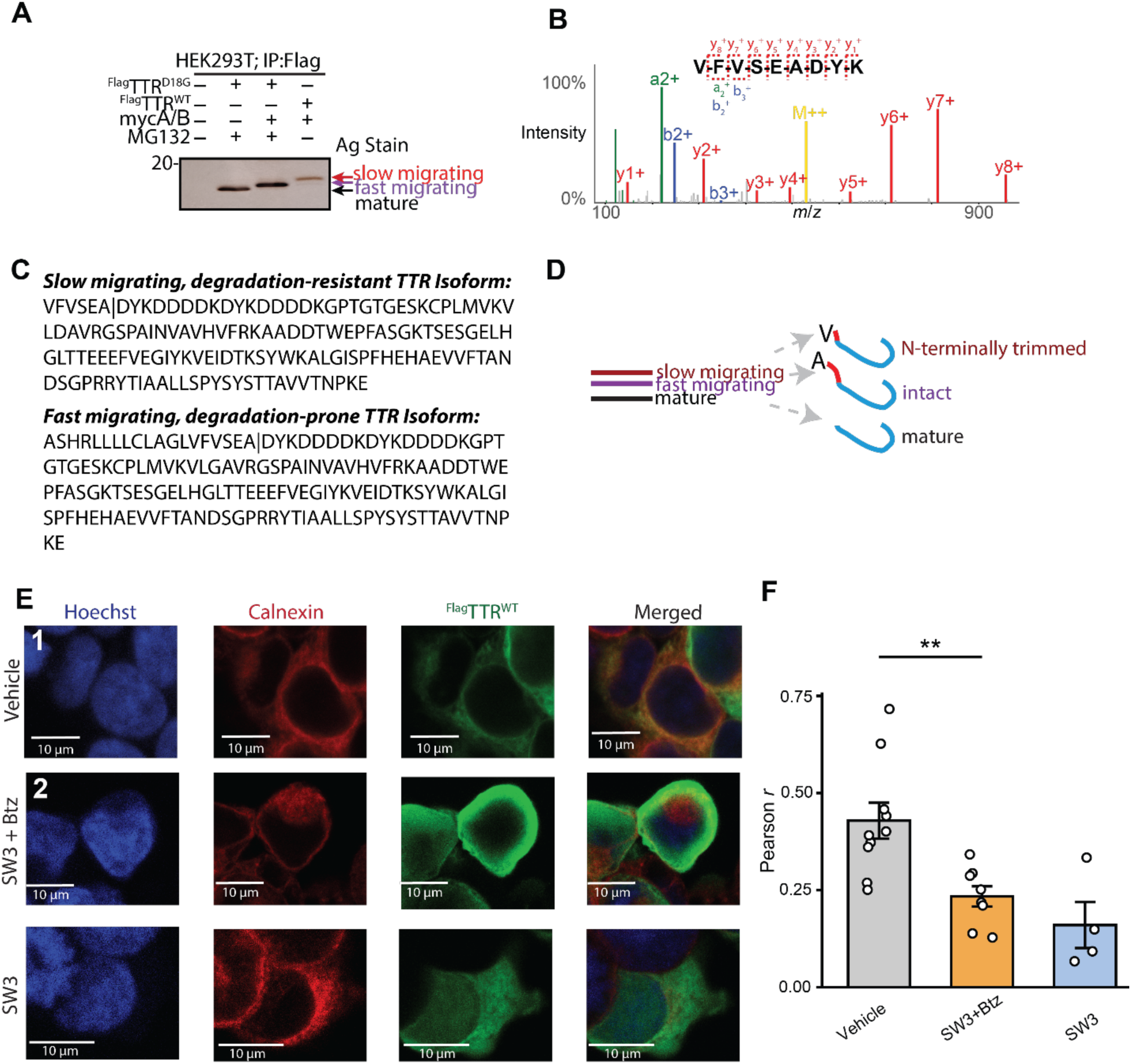
Cytosolic N-terminal Trimming Protects Mislocalized TTR^WT^ from Degradation. **A.** Silver stained SDS-PAGE-separated eluates from the immunoprecipitations prepared under the indicated conditions. **B**. Mass spectrum (MS^2^) and assignments for an N-terminal semi-tryptic peptide from mislocalized ^FLAG^TTR^WT^ that demonstrates N-terminal trimming to Valine. **C.** Sequences of TTR isoforms prepared as in **A**, based on mass spectrometric analysis. **D**. Schematic of TTR proteoforms. Illustration of the three major TTR species: (i) N-terminally trimmed cytosolic isoform (slow-migrating), (ii) intact precursors (fast-migrating), and (iii) ER-processed mature species. **E.** Confocal imaging of HEK293T cells expressing ^FLAG^TTR^WT^ treated with vehicle or SW3 + Btz, or their combination. Scale bars = 10 μm. Control samples lacking primary antibody have minimal background staining (**Supplemental Figure S3F**). **F.** Plot of the average Pearson’s *r* values between ^FLAG^TTR^WT^ and calnexin. Error bars represent S.E.M. n = 10 cells (vehicle), 8 cells (SW3+Btz; one cell excluded as a statistical outlier, Grubbs’ test, p = 0.045), and 4 cells (SW3). Hoechst channel contrast was enhanced 5% across the whole image for visualization; for the SW3-alone condition, the calnexin channel contrast was enhanced 2.5%.

The slow-migrating isoform has two main masses of similar intensity. One form has a mass of 16,539 Da, which corresponds to N-terminal trimming of the first 13 residues of ^FLAG^TTR^WT^ within the signal sequence (**Figure 3B-C**), and subsequent acetylation of the exposed N-terminal leucine. The other has a mass of 16,384 Da, representing trimming of 14 residues leaving an N-terminal valine. Shotgun proteomics of the tryptically digested protein confirmed the peptide sequences with MS2 spectra, where the cleavage in the signal sequence was at the valine (**Figure 3B-C**). The slow-migrating band will from now on be referred to as the N-terminally trimmed (see key on **Figure 3D**). We were surprised to see that the larger species migrates faster, and considered that this might be driven by the highly charged FLAG tag C-terminal to the signal sequence. We expressed a TTR^WT^ construct lacking the FLAG tag and observed the same two immature isoforms upon mycA/B treatment, but with the migration order (based on response to proteasome inhibitor treatment) now reversed and a slightly larger yield of the intact isoform (**Supplemental Figure S3B**).

There are two ways to interpret the partitioning of TTR between the degradation-resistant N-terminally trimmed form and the degradation-sensitive intact form. It could be that N-terminal trimming defines the triaging decision, with protein left intact now committed towards UPS. Alternatively, protein that evades UPS for an adequate period of time could be redirected for N-terminal trimming; in this case, triage does not depend on proteolysis. MG132 treatment robustly restores the intact isoform, without leading to an increase in the N-terminally trimmed form (**Figure 3A**), inconsistent with the latter interpretation. Hence, N-terminal trimming is likely the determinative step that partitions mislocalized protein between fates.

Because MycA/B and SW3 inhibit ER import, it is to be expected that the N-terminal trimming takes place in the cytosol. Alternatively, we considered the possibility that the trimmed population represents protein that evades Sec61 inhibition to enter the ER, is then trimmed in the ER in lieu of canonical signal sequence cleavage, and then re-enters the cytosol through retrotranslocation. To test this possibility, we tried two approaches at glycosylation-based reporters of ER entry, including an enhanced cryptic TTR sequon (T96F; **Supplemental Figure S3C**)^54^ and a previously described C-terminal NVT sequon (**Supplementary Figure S3D-E**).^55^ However, neither construct yielded detectable or interpretable glycosylation under these conditions, precluding definitive localization by this approach. Hence, we instead considered whether inhibiting ERAD would increase the yield of N-terminally trimmed TTR during Sec61 inhibition.

To test this, we treated cells with LS102, an inhibitor of HRD1 activity that acts by blocking its E3 ligase activity and consequent autoubiquitination.^56,57^ If the N-terminal trimming occurs in the ER and the trimmed product retrotranslocates to the cytosol, then blocking ERAD will result in more N-terminally trimmed isoform. The intact ^FLAG^TTR^WT^ isoform was partially rescued by LS102 and SW3 co-treatment relative to SW3 alone, but this co-treatment failed to increase the abundance of the N-terminally trimmed isoform (**Supplemental Figure S3G**). This rescue of the intact form could alternatively reflect inhibition of an off-target E3 ligase rather than HRD1 itself. Since HRD1 inhibition under Sec61 inhibition does not lead to an increase in the N-terminally trimmed form, it is unlikely that N-terminal trimming occurs in the ER.

We further used confocal immunofluorescence to define the cellular localization of TTR during Sec61 inhibition. Misfolded proteins that evade degradation can be segregated to specific locations within the cell such as inclusion bodies and stress granules.^59^ We considered that mislocalized TTR might be found distributed diffusely throughout the cytosol, or could be segregated into such compartments. Using confocal immunofluorescence, we examined how inhibition of Sec61 and/or the proteasome influence the cellular distribution of TTR. In vehicle-treated cells, ^FLAG^TTR^WT^ colocalizes with the ER resident protein calnexin (**Figure 3G row 1 and Figure 3H**. Under Sec61 inhibition and Btz treatment, which causes an accumulation of both intact and N-terminally trimmed forms (**Figure 2**), ^FLAG^TTR^WT^ signal remains distinct from the calnexin signal (**Figure 3G row 2 and Figure 3H**). Although the mislocalized protein population is no longer associated with the ER, we do not see evidence of puncta.

### Degradation Pathways of Mislocalized TTR Isoforms

To identify which cytosolic degradation pathways clear mislocalized TTR, we used chemical inhibitors targeting the two major pathways: the ubiquitin proteasome system and the autophagy lysosome system. Chloroquine (CQ), a lysosomotropic agent, broadly inhibits the acidification of cellular organelles. Bafilomycin A1, an inhibitor of V-ATPase, disrupts lysosomal acidification and autophagosome-lysosome fusion. SAR405, a PIK3C3 kinase inhibitor, inhibits conversion of LC3-I to LC3-II and prevents autophagosome formation.^60–62^ We treated HEK293T cells expressing ^FLAG^TTR^WT^ with mycA/B and each of these agents to assess their effects on the accumulation of mislocalized TTR isoforms (**Figures 4A,B**). To confirm the activity of the autophagy inhibitors, we monitored LC3 processing by immunoblot, assessing the conversion of LC3-I to LC3-II and its accumulation under autophagic flux inhibition (**Supplemental Figure S4A**). As observed above (**Figure 2A**), proteasomal inhibition (MG132) leads to increased levels of the fully intact TTR isoform. CQ treatment significantly elevates this intact TTR isoform, suggesting that mislocalized TTR is targeted for lysosomal degradation in addition to proteasomal degradation (**Figure 4A, lane 4**). Bafilomycin and SAR405 also partially restored the degradation-prone fully intact band, though to a lesser extent than CQ (**Figure 4A, lanes 5 and 6**). The N-terminally trimmed TTR isoform only slightly accumulates in response to each treatment, reflecting its resistance to degradation. Similar results were obtained for TTR^WT^ without a FLAG tag, indicating that the affinity tag does not affect the phenotype (**Supplemental Figure S4B**).

**Figure 4:**
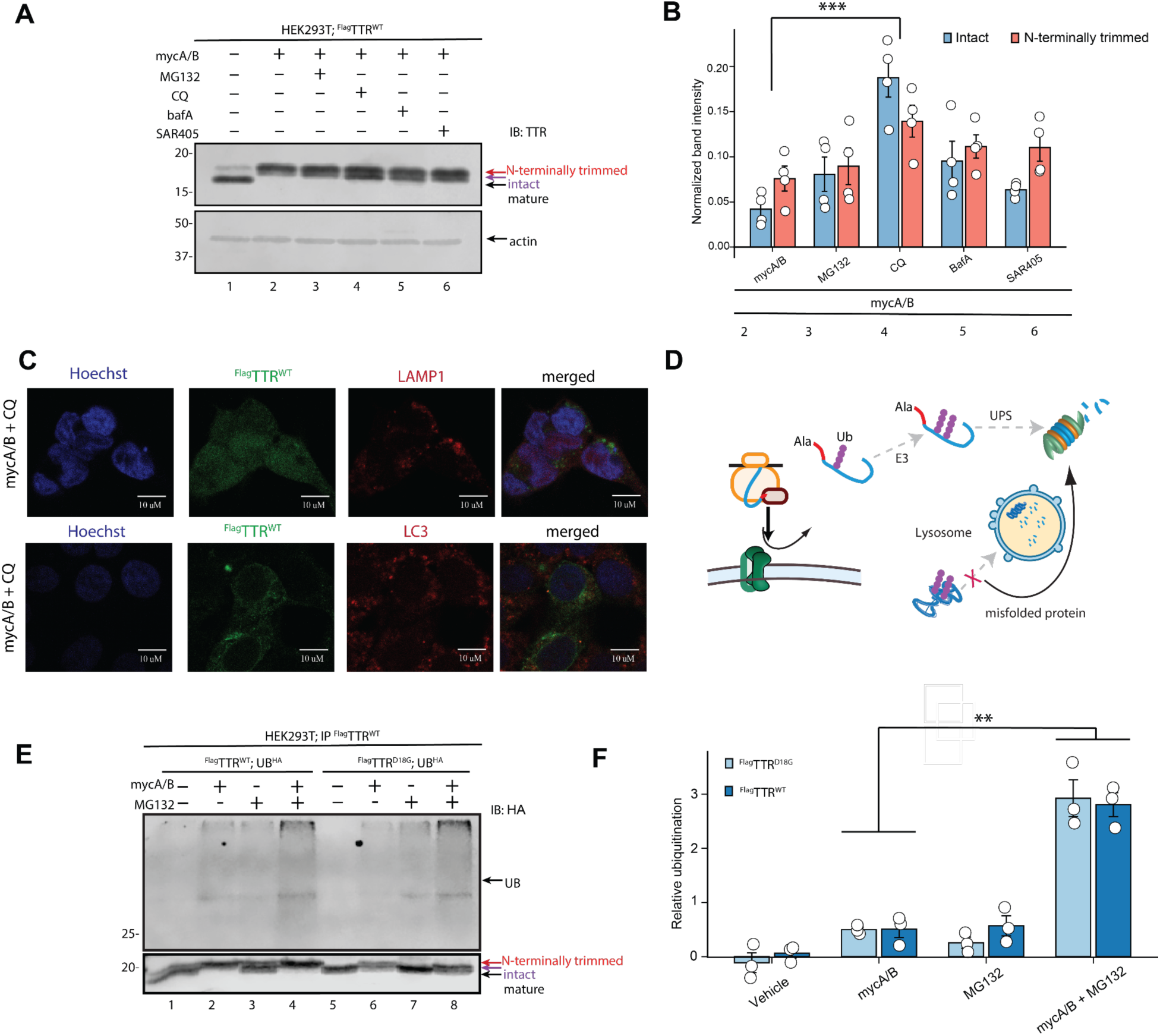
Degradation of mislocalized TTR isoforms is dependent on ubiquitin, the proteasome, and autophagy pathways. **A.** Representative immunoblot analysis of lysates from HEK293T Cells overexpressing ^FLAG^TTR^WT^. Cells were treated for 16 h with 25 nM mycA/B, 1µM MG132, 1µM SAR405, or 100µM chloroquine (CQ) or 100nM bafilomycin (BafA). Cells were then harvested and lysed. **B.** Quantification of immunoblots from HEK293T cell lysates expressing ^FLAG^TTR^WT^ and treated with indicated degradation inhibitors *** p< 0.01 by Student’s *t* test with Bonferroni correction. Grubbs’ test identified the highest BafA band intensity (13.622) as a significant outlier (*G* = 1.496, *p* = 0.0057). **C.** Confocal immunofluorescence of ^FLAG^TTR^WT^ in HEK293T cells treated with mycA/B + CQ. LAMP1^lysosomes^ (left, green) appear enlarged and clustered upon CQ treatment, but TTR A/B (magenta) remains diffuse with no detectable colocalization. Similarly, LC3^autophagosomal^ structures (right, red) show no spatial overlap with TTR, despite robust LC3 puncta accumulation. Pearson’s *r* = −0.30, Manders’ *tM*2 = 0.20; Costes *P* = 1.00; *n* = 20 cells. Hoechst marks nuclei in blue. Scale bars: 20 μm. **D.** Schematic illustrating how Sec61 translocon inhibition produces cytosolic TTR that is primarily degraded by the ubiquitin–proteasome system. Lysosomal blockade by CQ may redirect other misfolded clients toward the proteasome, reducing proteasomal flux and indirectly stabilizing mislocalized TTR. Lack of LC3 or LAMP1 colocalization suggests that TTR is not a macroautophagy cargo under these conditions. **E.** Representative immunoblot from HEK293T cell lysates co-expressing Ub-HA and either ^FLAG^TTR^WT^ or ^FLAG^TTR^D18G^ (25 nM mycA/B 16 h; 1 µM MG132 16 h). **F.** Quantification of immunoblots from 3E *n* = 3. Significance codes: *** p < 0.001; ** p < 0.01; * p < 0.05. Error bars represent S.E.M.

Since CQ robustly rescues the intact TTR isoform in our immunoblots, we next asked whether mislocalized TTR is routed to autophagosomes and/or to lysosomes for degradation. We performed confocal microscopy for TTR together with markers for either the lysosome (LAMP1) or autophagosome (LC3). Under mycA/B and CQ treatment, LAMP1 forms enlarged, ring-like structures consistent with lysosomal swelling (**Figure 4C**). However, across multiple fields and z-stacks, we observed no colocalization of LC3 or LAMP1 with ^FLAG^TTR^WT^. Under mycA/B + MG132 + CQ conditions, ^FLAG^TTR^D18G^ also did not colocalize with LC3-positive puncta and upper optical sections suggested localization with cell periphery, without forming discrete puncta (**Supplemental Figures S4C–D**). Under sec61 inhibition and proteasome inhibitor, ^FLAG^TTR^D18G^ has a localization pattern similar to ^FLAG^TTR^WT^, with both diffuse cytosolic staining and peripheral signal suggestive of cell surface localization (**Supplemental Figure S4D-E**).

Thus, although CQ prevents the degradation of mislocalized TTR, our imaging suggests that this effect is indirect and does not result from the direct sequestration of TTR within autophagosomes or its delivery to lysosomes. The autophagy-lysosomal system has been shown to serve as a compensatory degradation pathway during proteasomal saturation.^63^ If lysosomal blockade redistributes degradative load to UPS, that could reduce proteasomal capacity to degrade mislocalized TTR (**Figure 4D**).

Ubiquitin chains linked through the same site on each monomer play key roles in cellular processes: K48-linked chains target proteins for degradation, while M1- and K63-linked chains are involved in DNA repair, NF-κB signaling, and autophagy.^64^ We hypothesized that TTR ubiquitination would increase under conditions of ER import inhibition and proteasomal inhibition, as the degradation-prone intact form accumulates in the cytosol. To evaluate ubiquitination of mislocalized TTR, we co-overexpressed ^FLAG^TTR^WT^ or ^FLAG^TTR^D18G^ with Ub-HA in HEK239T cells. Cells were lysed in the presence of the deubiquitinase inhibitor NEM, ^FLAG^TTR immunoprecipitated with anti-FLAG beads, and ubiquitination evaluated by immunoblotting. Little ^FLAG^TTR ubiquitination is observed in the absence of Sec61 translocon or proteasomal inhibition (**Figure 4E lanes 1 and 5, and Figure 4F**). MycA/B treatment alone increases ubiquitination relative to vehicle control (**Figure 4E, lanes 2 and 6**), indicating that mislocalization due to impaired ER translocation enhances recognition and tagging by ubiquitin ligases. Among all conditions tested, co-treatment with both MG132 and mycA/B (**Figure 4E, lanes 4 and 8**) resulted in far higher levels of ubiquitinated ^FLAG^TTR, with similar relative ubiquitination of both the WT and D18G TTR variants. Ubiquitination of mislocalized ^FLAG^TTR is primarily serving as a signal for proteasomal targeting.

### Mislocalized TTR Aggregation

When ER import is disrupted, for example by ER stress, secretory protein precursors remain in the cytosol, where they are prone to misfolding and aggregation. Similarly, secretory protein translocation inhibition by Hspa13 dysregulation causes accumulation and aggregation of TTR.^12^ To test whether TTR mislocalized via Sec61 translocon inhibition can fold, we treated HEK293T cells overexpressing ^FLAG^TTR^WT^ or ^FLAG^TTR^D18G^ with mycA/B and Tafamidis (Taf, 10 μM), which stabilizes TTR in its native tetrameric state and prevents dissociation.^65,66^ If nascent mislocalized TTR can fold and achieve the tetrameric state, Taf binding should shift the equilibrium away from the unfolded monomer, and protect it from degradation. Taf treatment affects neither mislocalized TTR degradation nor the abundance of the mislocalized TTR isoforms, indicating that cytosolic TTR is not in equilibrium with the tetrameric state and is misfolded in the cytosol (**Supplemental Figure S5A**).

Since mislocalized TTR does not reach the tetrameric state, we hypothesized that it would aggregate upon Sec61 translocon inhibition. We performed ultracentrifugation to separate insoluble aggregates from soluble protein in lysates of cells expressing ^FLAG^TTR^WT^ under different treatment conditions (mycA/B, MG132, and/or CQ; **Figure 5A**). MycA/B treatment, which leaves behind the degradation-resistant, N-terminally trimmed isoform, causes only a minor increase in insoluble TTR aggregates (**Figure 5B, lane 4**). In contrast, cotreatment with mycA/B and MG132 stabilizes the mislocalized intact isoform, leading to a dramatic increase in insoluble TTR aggregate levels (FC =5.39 ± 2.75 vs. vehicle; FC =2.71 ± 0.99 vs. mycA/B alone; **Figure 5B**). CQ and mycA/B treatment produced a similar effect, corresponding with the rescue of the same intact isoform (**Figure 5B, lane 8**). MycA/B with either or both degradation inhibitors significantly increases the aggregate compared to mycA/B alone (**Figure 5C**). These results suggest that retention of the hydrophobic signal sequence on the intact TTR isoform increases misfolding, thereby promoting aggregation, and that its removal via N-terminal trimming reduces this aggregation propensity. This is consistent with our Taf results (**Supplemental Figure S5A**): Taf treatment affects neither mislocalized TTR degradation nor the abundance of either mislocalized TTR isoform, indicating that cytosolic TTR is not in equilibrium with the tetrameric state and is misfolded in the cytosol. To assess differences in chaperone binding between the intact and trimmed forms of mislocalized TTR, we performed immunoprecipitation-mass spectrometry on treated HEK293T cells overexpressing ^FLAG^TTR^WT^ with mycA/B in the presence or absence of Btz. Cells were treated with the reversible cell-penetrable crosslinker DSP to preserve interactions, lysed, and TTR was immunoprecipitated. The crosslinking was reversed, and proteins digested to peptides and labeled with isobaric TMT tags for bottom-up mass spectrometric analysis of interactor enrichment. Protein levels were then normalized to TTR levels for each sample. Btz treatment increases the amount of the intact TTR, while leaving the N-terminally trimmed population unaffected, so interactors relatively enriched under this condition will be preferential intact TTR interactors (**Supplemental Figure S5B**). There was an enrichment for chaperone interactors with the intact TTR isoform, which further validates its misfolded state. These findings underscore the protective role of preferential degradation in maintaining cytosolic proteostasis.

**Figure 5:**
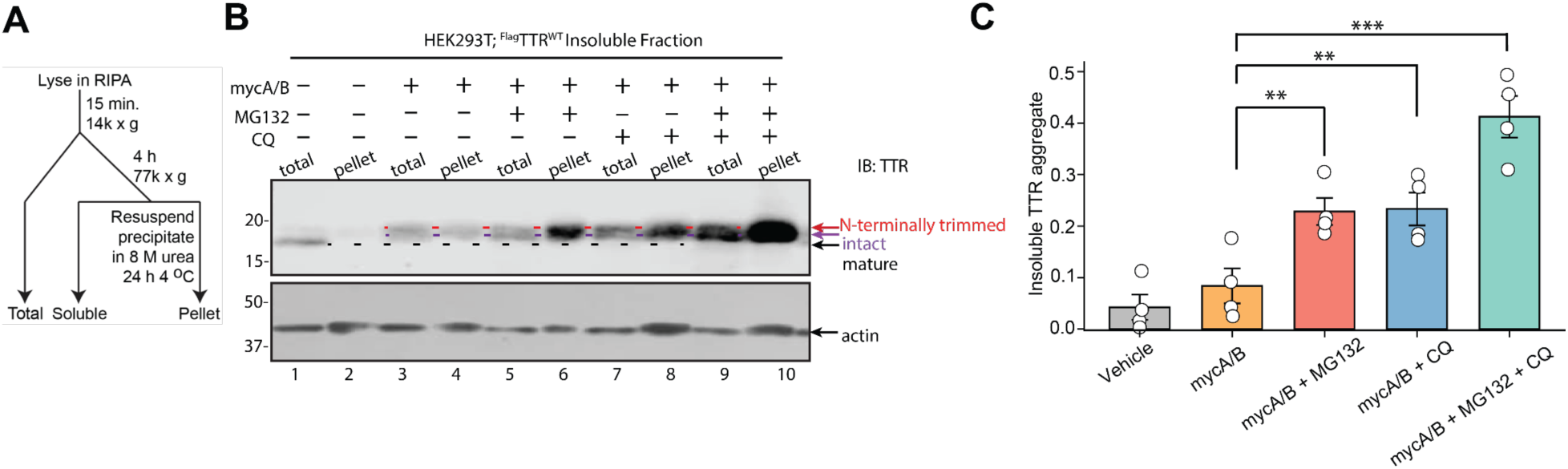
Intact Mislocalized TTR forms insoluble aggregates. **A.** The schematic details how the insoluble protein is isolated. **B.** Representative immunoblot of SDS-PAGE-separated lysates as total protein (total) and insoluble protein fraction (pellet) from HEK239T cells overexpressing ^FLAG^TTR^WT^ and treated for 16 h as indicated (25 nM mycA/B; 1 µM MG132; 100 µM CQ). Actin blotting is shown to demonstrate equivalent loading of total lysates. **C.** Quantification of figure 4A. Post-hoc Tukey’s HSD test (ANOVA F = 20.94 < Fcrit = 3.05; n = 4). Significance codes: *** p < 0.001; ** p < 0.01; * p < 0.05. Bars represent mean ± SEM; circles represent individual biological replicates (n = 4).

We next hypothesized that degradation of destabilized variants might mitigate inherent aggregation propensity to lower the overall load of aggregate protein associated with mislocalization. To test this, we compared overexpressed ^FLAG^TTR^WT^ (thermodynamically stable), ^FLAG^TTR^L55P^ (metastable), and ^FLAG^TTR^D18G^ (highly destabilized) in HEK293T cells using the same ultracentrifugation assay described in **Figure 5A**. Under vehicle conditions, all three of the TTR variants form little to no aggregate (**Supplemental Figure S5C and Supplementary Figure 5D**). Upon proteasome inhibition (**Supplemental Figure S5D, lanes 7–9**), TTR^D18G^ has a similar accumulation of aggregates as WT, consistent with rescue of degraded protein. ^FLAG^TTR^L55P^ remained the least aggregated, despite having thermodynamic and kinetic stability intermediate between D18G and WT. This suggests that the inherent amyloidogenicity of the mature proteins does not primarily drive the aggregation propensity of the mislocalized proteins.

### Role of Thermodynamic Stability in Mislocalizing Protein Quality Control

We found above that the cell differentiates between the thermodynamically stable ^FLAG^TTR^WT^ and destabilized ^FLAG^TTR^D18G^, leading to the former being partially trimmed to lower both aggregation and degradation propensity, while the latter is primarily ubiquitinated and degraded. There are two major biophysical parameters differentiating these two variants. ^FLAG^TTR^D18G^ is unable to form tetramers, and hence lacks the kinetic stability of ^FLAG^TTR^WT^. ^FLAG^TTR^D18G^ is also highly thermodynamically destabilized, with misfolded states strongly preferred over the native TTR folding. To better understand the factors governing triaging of misfolded TTR, we took advantage of the extensive biophysical characterization of TTR variants.^37^ Using this framework, we tested whether the efficiency of N-terminal trimming depends on the thermodynamic or kinetic stability of TTR. We analyzed variants spanning a range of thermodynamic (C_m_; the concentration of urea needed to reach the midpoint of denaturation) and kinetic stability (t_½_; the rate of dissociation from tetramer to monomer) and quantified their N-terminal processing (**Figure 6A**). When we compared the accumulation of intact TTR under Sec61 and proteasome inhibition with the thermodynamic stability of each variant, we found that lower thermodynamic stability was generally associated with greater accumulation of the intact species (**Figure 6B**). In contrast, the relationship between accumulation of intact TTR with kinetic stability is less consistent. This distinction is mostly driven by the V30M variant, which deviates from the kinetic stability trend. Specifically, V30M has kinetic stability comparable to WT but similar thermodynamic stability to A25T and D18G. This result suggests that thermodynamic stability is a more consistent predictor of N-terminal trimming efficiency than kinetic stability, consistent with the inability of the kinetic stabilizer Taf to influence trimming (**Figure S5A**).

**Figure 6:**
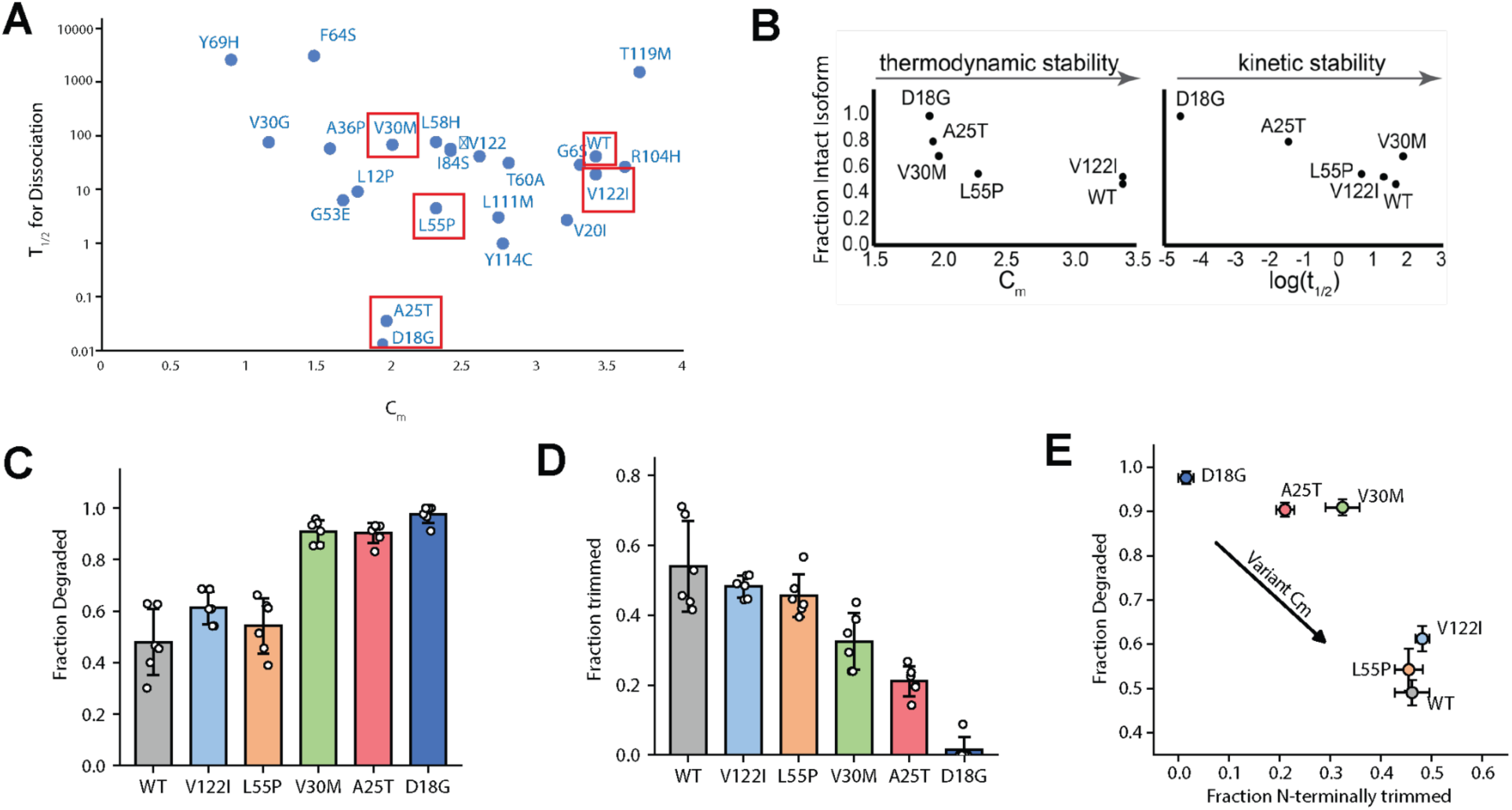
Cytosolic Quality Control of Thermodynamic Stability of TTR. **A.** Thermodynamic (Cm) and kinetic (t1/2 for tetramer dissociation) stability parameters of TTR variants, adapted from Sekijima et al. Variants used in this study are highlighted. **B.**. Comparison between energetic stability and the fraction of intact intermediate TTR isoform detected under mislocalization. The fraction of intact isoform was quantified as (intact intermediate) / (intact + trimmed) under mycA/B+ MG132 conditions and plotted against published Cm or t1/2. **C.** Steady state degradation yield of mislocalized TTR variants. Degradation was calculated as 1 − (signal intensity under mycA/B / signal intensity under mycA/B + MG132 + CQ), representing the proportion of intact TTR removed by the proteasome. **D.** Extent of N-terminal trimming for each TTR variant. The fraction of trimmed isoform was calculated as (trimmed) / (trimmed + intact) under mycA/B + MG132 treatment. Stable variants accumulate more trimmed products. Bars show mean ± S.E.M; open circles represent individual biological replicates (n = 6 per variant, except D18G mycA/B-only condition). **E.** Comparison between fraction of intact intermediate isoform and fraction degraded for each variant. Thermodynamically destabilized variants (D18G, A25T, V30M) cluster toward high degradation and low trimming; stable variants (WT, V122I, L55P) exhibit the opposite trend.

We next quantified how efficiently each variant is degraded upon mislocalization for the following variants: ^FLAG^TTR^WT^, ^FLAG^TTR^V122I^, ^FLAG^TTR^L55P^, ^FLAG^TTR^V30M^, ^FLAG^TTR^A25T^, and ^FLAG^TTR^D18G^ (**Figure 6C**). The degradation yield was calculated as the amount of mislocalized TTR protected by the presence of degradation inhibitors, as a fraction of total mislocalized TTR. The more thermodynamically stable ^FLAG^TTR^WT^, ^FLAG^TTR^V122I^ and ^FLAG^TTR^L55P^ are degraded less efficiently than highly thermodynamically destabilized ^FLAG^TTR^V30M^, ^FLAG^TTR^A25T^ and ^FLAG^TTR^D18G^ (**Figure 6C**). Each variant was overexpressed in HEK293T cells under conditions of mycA/B treatment, alongside MG132 and CQ treatment, and the relative trimming yield determined as the fraction of trimmed TTR against total immature TTR (**Figure 6D**). When combining the steady-state degradation, fraction of intact isoform, and thermodynamic stability are examined together (**Figure 6E**), a model emerges: thermodynamically unstable TTR variants are recognized and eliminated by cytosolic QC pathways, whereas more stable variants can evade degradation, undergo N-terminal trimming, and persist in the cytosol. For each of these variants, proteasome or lysosome inhibition does not increase the amount of N-terminally trimmed isoform, similar to what we observed for ^FLAG^TTR^WT^ and ^FLAG^TTR^D18G^ (**Figures 2B and 4B**), and consistent with trimming occurring upstream of degradation rather than in competition with it.

## DISCUSSION

Canonically, quality control of mislocalized secretory proteins involves their degradation. We find that thermodynamically stable mislocalized secretory proteins stay persistent in the cytosol and escape degradation and aggregation in the cytosol through an N-terminal trimming. This N-terminus trimming pathway is distinct from other ER quality control mechanisms, where destabilized proteins are marked for degradation through ERAD by either mannose trimming or ubiquitination post-retrotranslocation.^67,68^ We find that both persistent intact and N-terminally trimmed TTR under Sec61 inhibition do not co-localize with an ER resident protein, indicating a cytosolic quality control mechanism. Unlike degradation-only models of cytosolic quality control, this N-terminal trimming pathway discriminates proteins based on their intrinsic energetic parameters when proteasomes are overwhelmed and NRF1 compensation is insufficient.

It has been proposed that stress-induced ER preQC can be mediated by Derlin upregulation at the ER membrane downstream of the IRE1 arm of the UPR.^13,69^ An alternative proposal is that ER preQC is not mediated by a decrease in Sec61 translocon activity, but rather by PERK-dependent downregulation of SRP14, and consequent inefficient SRP targeting.^14^ In the IRE1-dependent model for preQC, nascent mislocalized proteins are released into the cytosol at the rough ER, while in the PERK-dependent model they are not necessarily generated close to the ER. We see N-terminal trimming subsequent to both Tg-induced ER preQC (**Figure 1D**) as well as direct Sec61 translocon inhibition. If ER preQC is primarily driven by a failure to engage SRP, then the N-terminal trimming mechanism will have to act similarly for secretory protein synthesized at blocked Sec61, and secretory protein synthesized at ribosomes failing to engage SRP14.

Cytosolic N-terminal trimming functions as cellular protection under secretory load that challenges degradation capacity, which could be particularly important for high secretory load tissues such as hepatocytes, lymphocytes, and pancreatic beta cells, where Sec61 translocon and the proteasome may be more readily overwhelmed. Our evidence on Tg-induced mislocalization could expand to other environmental toxins that disrupt SERCA pumps, such as cadmium^70^, bisphenyl^71^, alkylphenols, as well as other sources of ER stress, which may also lead to mislocalization and N-terminal trimming. Generally, mislocalization of proteins has been reported in diabetes,^22,23,34,1^ neurodegeneration,^15^ and pathogenesis,^22,23,34^ and this trimming might be important for maintaining cytosolic proteostasis in those contexts as well. We found that N-terminal trimming selects TTR substrates based on thermodynamic stability. Because the relevant mutations lie far past the signal sequence and none of the TTR variants fold properly in the cytosol, we still need to determine the driving factors of this recognition.

Cytosolic N-terminal trimming emerges as a protective adaptation during proteostasis stress, selectively acting on thermodynamically stable secretory proteins. These findings expand our understanding of how the cytosol triages mislocalized secretory proteins and reveals a new layer of cytosolic quality control.

## MATERIALS AND METHODS

### Plasmid DNA

Wild-type ^FLAG^TTR and its variants (N-terminal FLAG) and Ub-HA have been previously described.^42^ ^FLAG^A1AT^WT^ with its native signal sequence and ^HA^A1AT^NHK^ were provided by Lars Plate (Vanderbilt University). Preprolactin was a custom vector by Genscript. Sequences can be found in Supplemental Information.

### Cell culture

Human embryonic kidney cell line 293 with T antigen (HEK293T) were grown in Dulbecco’s Modification of Eagle’s Medium (DMEM; Corning 10-013-CV) containing 10% fetal bovine serum, 1% L-Glutamine and 1% Penicillin-Streptomycin. HEPG2 cell lines were grown in Eagle’s Minimum Essential Medium (EMEM; MT10009CV) containing 10% fetal bovine serum and 1% Penicillin-Streptomycin. Cells were transfected at 20-60% confluence while being attached to the plates. All transfection performed in the study were Ca_3_(PO_4_)_2_ transfection. 250 mM CaCl_2_ containing DNA plasmid was vortexed with drop-wise addition of 2X HEPES-buffered saline phosphate solution (2X HBS). The resultant transfection solution was added drop-wise to the cells with fresh replenishment of media after 12-16 hours. To enhance the adherence of HEK293T cells during drug treatments, cells were reseeded on poly-D-lysine treated plates. To inhibit ER translocation, we treated the transfected cells with 25 nM mycolactone (mycA/B) or 50 nM SW3 for 16 hours. To prevent degradation of mislocalized proteins in the cytosol, 1 μM MG132, 1μM bortezomib (Btz) and/or 100 μM chloroquine (CQ) was added to the cells for 16 hours. The negative control received the same volume of DMSO vehicle.

### Cell lysis

Cells were lysed in RIPA buffer (50 mM Tris, 15 mM NaCl, 1% Triton X100, 0.5% deoxycholate and 0.1% sodium dodecyl sulfate (SDS)) containing 1X EDTA-free protease inhibitor cocktail (PI) (Roche) for at least 15 minutes on ice. The lysate was collected from the supernatant of a 21,100 g spin for 15 minutes. Protein was quantified from Bradford assay (Bio-rad) by spectrophotometry (Agilent Cary 50 and 60).

### Immunoblot sample preparation

The proteins from different samples were normalized and brought to the same volume by RIPA buffer. The 6X Laemmli loading dye was added to a final concentration of 1X and 16.7 mM dithothreitol was also included in the samples. To denature and reduce the proteins, the samples were boiled at 100°C for 5 minutes or 20 minutes as indicated. Proteins were loaded on freshly prepared SDS-polyacrylamide gel electrophoresis (SDS-PAGE) gels cast in the Bio-Rad Mini-PROTEAN system. The samples were run through the stacking gel and the resolving gel at 60 V and 170 V, respectively.

The gels, filter paper and nitrocellulose membrane were immersed in Towbin’s buffer, consisting of 25 mM Tris, 192 mM glycine, 20% methanol (per Bio-Rad transfer buffer recipes). The semi-dry transfer of the sandwich was conducted in a Bio-Rad Trans-Blot Turbo for 30-60 minutes at 25 volts and 1 A. The quality of the transfer was confirmed by Ponceau S assay. The membranes were then blocked with 5% milk in Tris-buffered saline (TBS) for 40-60 minutes. Blots were rinsed well with TBS buffer with 0.1% Tween (TBST) for several rounds. The proteins were then probed with primary antibody diluted in 5% bovine serum albumin (BSA), 0.1% NaN_3_ in TBS for at least two hours. Primary antibodies included: rabbit anti-TTR (DAKO; 1:1,000), mouse anti-actin (Proteintech; 1:2,000), mouse anti-FLAG (Sigma; 1:1,000), rabbit anti-LC3 (Proteintech; 1:1,000), or rabbit anti-HA (Proteintech;1:1,000).The membranes were then rinsed with TBST and probed with IR-dye conjugated secondary antibody (Li-COR) targeting the primary host such as rabbit or mouse for 30 minutes. The secondary antibody was diluted in 1:10,000 ratio in the blocking buffer. The membranes were repeatedly rinsed with TBST, followed by a rinse in TBS and water. After all steps of blotting, the membranes were imaged with a Li-Cor Odyssey Fc, using 700 nm and 800 nm filters and exposure times of 10 minutes each. The images had contrast adjusted image-wide with Li-Cor’s Image Studio, with densitometric quantification performed if necessary.

### EC50 analysis

Band intensities from immunoblots of FLAG–tagged TTR overexpressed in HEK293T cells were quantified in Li-Cor Image Studio. For each lane, a freehand ROI was drawn around the total TTR signal (immature + mature), and a second ROI was drawn around the immature TTR band. The fraction of immature TTR represents the immature over the total TTR signal. These values were plotted against inhibitor concentration and fit to a four-parameter log-logistic model using the drc (analysis of dose-response curves) package in R to obtain EC₅₀ estimates.

### Ultracentrifugation assay

Cell lysates after normalization were balanced in weight with RIPA in Beckman-Coulter polypropylene ultracentrifuge tubes (cat# 357448) and spun down in a Beckman-Coulter Optima Max-XP ultracentrifuge, using a TLA-55 rotor. Samples were spun at 77,000 x g for four hours, at 4°C and under vacuum. Pellets formed from ultracentrifuge were rinsed four times with RIPA and incubated for 3-4 days with 8 M urea at 4°C. The solubilized pellets were diluted to 2 M urea with RIPA and prepared for immunoblot by addition of reducing Laemmli and boiling for 5 minutes.

### Immunofluorescence

HEK293T cells transiently expressing FLAG-tagged TTR were reseeded onto poly-D-lysine–coated glass-bottom 24-well plates (Chemglass Life Sciences) and treated 4 h later with 25 nM mycA/B, 1 μM MG132, bafilomycin, or vehicle for 16 h as indicated. Cells were fixed in 10% formalin in fixation buffer (60 mM PIPES, 25 mM HEPES, 10 mM EGTA, 2 mM MgCl₂, 0.12 mM sucrose, pH 7.4) for 15 min at room temperature, washed with DPBS, and permeabilized in pre-chilled methanol at −20 °C for 5 min. After three DPBS washes, samples were blocked overnight in 3% BSA/0.1% NaN₃ in DPBS. Primary antibodies were applied overnight in 1% BSA/0.1% NaN₃, including mouse anti-FLAG (1:250), mouse anti-LAMP1 (DSHB; 1:200), rabbit LC3 (Proteintech; 1:250) or Coralite Plus-647-conjugated Calnexin Rabbit PolyAb. After four DPBS washes, Alexa Fluor secondary antibodies were applied for 4 h: goat anti-rabbit IgG Fab2 Alexa Fluor 594 (1:1000) for organelle markers and alpaca anti-mouse IgG1 Fc Alexa Fluor 488 (1:1000) for FLAG. For experiments requiring multiple primaries from different hosts, sequential staining and wash steps were performed. Nuclei were stained with Hoechst 33258 (TOCRIS Bioscience; 1:15,000) for 10 min prior to final washes. Imaging was performed using a Zeiss LSM 880 confocal microscope with a 100×/1.46 NA oil objective. Alexa Fluor 594, Alexa Fluor 488, and Hoechst were imaged using 594-, 488-, and 355-nm excitation, respectively, with corresponding emission windows (594: 599–633 nm, 488: 500–535 nm, Hoechst: 381–480 nm). All imaging parameters were kept constant within each experiment. Representative cells were selected using Region of Interest (ROI) tools to isolate the cytoplasm and exclude extracellular background. Prior to analysis, images underwent background subtraction in both channels to ensure signal integrity. The Costes automated thresholding method was applied to calculate Pearson’s Correlation Coefficient (r) and Manders’ Split Coefficients (tM1 and tM2). Statistical significance of the correlation was determined via the Costes Significance Test using 100 randomizations, with a P-value > 0.95 considered significant. All results were compiled and processed using a Fisher r-to-z transformation to determine the mean Pearson value across the dataset.

### RNAseq

RNAseq was performed by Plasmidsaurus. 200,000 HepG2 cells per sample subjected to indicated treatment conditions were removed using TrypLE protease solution, pelleted, and resuspended in RNAguard. Quality of the fastq files was assessed using FastQC v0.12.1. Reads were then quality filtered using fastp v0.24.0 with poly-X tail trimming, 3’ quality-based tail trimming, a minimum Phred quality score of 15, and a minimum length requirement of 50 bp. Quality-filtered reads were aligned to the reference genome using STAR aligner v2.7.11 with non-canonical splice junction removal and output of unmapped reads, followed by coordinate sorting using samtools v1.22.1. PCR and optical duplicates were removed using UMI-based deduplication with UMIcollapse v1.1.0. Alignment quality metrics, strand specificity, and read distribution across genomic features were assessed using RSeQC v5.0.4 and Qualimap v2.3, with results aggregated into a comprehensive quality control report using MultiQC v1.32. Gene-level expression quantification was performed using featureCounts (subread package v2.1.1) with strand-specific counting, multi-mapping read fractional assignment, exons and three prime UTR as the feature identifiers, and grouped by gene_id. Final gene counts were annotated with gene biotype and other metadata extracted from the reference GTF file. Sample-sample correlations for sample-sample heatmap and PCA were calculated on normalized counts (TMM, trimmed mean of M-values) using Pearson correlation. Differential expression was done with edgeR v4.0.16 using standard practice including filtering for low-expressed genes with edgeR::filterByExprwith default values. Functional enrichment, when available, is performed using gene set enrichment analysis with gseapy v0.12 using the MSigDB Hallmark gene set. Analysis was restricted to protein-coding genes. Low-expression genes were removed by requiring a mean CPM ≥ 1 across all samples within each dataset, retaining 12,175 of 20,120 protein-coding genes for the SW3 dataset and 11,951 of 16,826 protein-coding genes for the mycA/B dataset. For each gene, fold change was calculated as the ratio of mean normalized expression in the treatment condition to mean normalized expression in vehicle (or, for WRR139 combination conditions, relative to the corresponding single-agent condition). Statistical significance was assessed using a two-sample t-test per gene (3 replicates per group), followed by multiple-testing correction using Storey’s q-value method, a modification of the Benjamini-Hochberg procedure. Genes were considered significantly differentially expressed at q < 0.05 and |log_2_FC| ≥ 1. We used Claude to edit and fix R code to visualize data. No AI was used for image processing, and we reviewed and edited the code as needed and take full responsibility for the plots that were generated. The RNA-sequencing data generated in this study have been deposited at Gene Expression Omnibus under accession GSE347460 and are available as of the date of publication.

### Peptide Mapping of Proteolytic Product of Mislocalized TTR

HEK293T cells were transfected with ^FLAG^TTR^D18G^ and treated with 25 nM mycA/B and 1 μM MG132 to obtain the majority of the intact band. Cells were then harvested and lysed in RIPA/PI. The lysates were precleared with Sepharose for 30 minutes at 4°C, and incubated with M2 anti-FLAG beads (Sigma) overnight at 4°C. The beads were then washed stringently with RIPA buffer for 4 times and eluted in Laemli buffer by boiling for 5 minutes. The eluate was precipitated by methanol-chloroform precipitation. Protein samples were brought up to 100–150 μL with H₂O. Methanol (400–450 μL) was added and the sample vortexed, followed by chloroform (100–150 μL) and a second vortex step. Water (300–450 μL) was then added, and the sample was vortexed thoroughly and centrifuged to induce phase separation. The upper aqueous layer was carefully aspirated without disturbing the protein pellet at the solvent interface. Methanol (500 μL) was added to the remaining sample, which was vortexed, centrifuged, and the supernatant aspirated, leaving no more than 20 μL of residual liquid. This methanol wash was repeated two to four additional times, depending on the extent of detergent contamination. Samples were centrifuged a final time to remove residual liquid from the tube walls, and the resulting pellet was air-dried in a fume hood for 0.5–2 hours. The air-dried protein pellet was resuspended in 9 M urea, reduced with 5 mM tris(2-carboxyethyl)phosphine for 30 minutes at room temperature, and alkylated with 10 mM iodoacetamide for 15 minutes at room temperature in the dark. The solution was diluted to 2 M urea by 50 mM Tris pH 8.0. The sample was digested with 0.5 ug trypsin overnight at 37°C overnight. The digestion was quenched by 0.5% formic acid and spun down at 21,100 x g for at least 30 minutes to remove debris.

The samples were preloaded on injected to a 2-cm trapping column containing C18 resin. Separation was achieved by applying 7-55% gradient of solvent B (80% ACN, 0.1% formic acid) at a flow rate of 500 nL/min. A voltage of 3.0 kV was applied for electrospray ionization and the inlet capillary was heated to 275°C. Data-dependent acquisition of MS/MS spectra with the LTQ-Orbitrap-Velos-Pro was performed with the following settings: MS/MS on the 10 most intense ions per precursor scan, 1 microscan, rejection of unassigned and charge state 1; dynamic exclusion repeat count 1, repeat duration, 30s; exclusion list size 500, and exclusion duration, 60s. The MS/MS spectra were obtained from CID fragmentation in the Ion trap of 38.0. Peptide identification was performed on Integrated Proteomics Pipeline IP2. Tandem mass spectra were extracted from the raw files using Rawconverter and were searched on a Uniprot human database with reversed sequences using the ProLucid algorithm.^73–75^ The search parameters included nontryptic peptides. Carbamidomethylation (+57.02146) of cysteine and tandem mass labeling (+229.1629) of lysine and N-terminus were set as static modification. Peptide candidates were filtered using DTASelect, with these parameters: -p 2 –y 0 –sfp 0.01 –DM 10. We require at least 2 identified peptides for a protein to be accepted. The peptides can be accepted regardless of enzyme specificity. The peptide level false positive rate was set to 0.01. We accepted peptides that have delta masses no more than 5 ppm.

### Crosslinking Mass Spec

HEK293T cells expressing FLAG-tagged TTR were treated for 16 h with 25 nM mycA/B together with vehicle or 1 µM bortezomib. Protein interactions were stabilized by adding the reversible, cell-permeable crosslinker DSP (1 mM, 30 min, RT), followed by quenching with final concentration 90 mM Tris-HCl (pH 7.5). Cells were lysed in RIPA lysis buffer supplemented with protease inhibitors, clarified by centrifugation, and ^FLAG^TTR complexes were isolated by anti-FLAG immunoprecipitation. Prior to IP, sepharose beads were used to remove nonspecific binding (4°C, 30 min). Crosslinks were reversed with DTT-containing elution buffer, and eluates were reduced, alkylated, and digested with trypsin as described above. Peptides were labeled with TMT-6plex isobaric reagents, quenched with ammonium bicarbonate, and pooled for multiplexed quantitative mass spectrometry. MS/MS analyses were performed using a two-dimensional LC/MS/MS setup on an LTQ Orbitrap Velos Pro hybrid mass spectrometer (Thermo) interfaced with an Easy-nLC 1000 (Thermo). Peptides were separated on an in-house packed biphasic MudPIT column built from fused-silica capillary (150 µm ID × 375 µm OD, Polymicro/Molex), packed with Luna® 5 µm SCX (100 Å, Phenomenex) followed by Aqua® 5 µm C18 (125 Å, Phenomenex). The mass spectrometer was operated in positive ion mode with a nanospray ionization (NSI) source (capillary temperature 275°C, source voltage 3.00 kV, source current 100 µA). Data-dependent MS/MS was performed on the top 10 most intense precursor ions per full scan, using higher-energy collisional dissociation (HCD) with an isolation width of 1.00 m/z, normalized collision energy of 38 (energy width 12.0, 3 collision energy steps), and a minimum signal threshold of 500.0. Dynamic exclusion was enabled (repeat count 1, repeat duration 30 s, exclusion duration 60 s, exclusion mass width ±1.50 Da, exclusion list size 500). MS/MS spectra were searched using FragPipe against a UniProt human reviewed proteome (UP000005640, dated 2024-04-09) with decoy sequences and common contaminants appended.^76^ Searches were performed using MSFragger with a precursor mass tolerance of 20 ppm and a fragment mass tolerance of 20 ppm, specifying trypsin as the digestion enzyme (cleavage at K/R, not before P) with full enzymatic specificity and up to 2 missed cleavages. Static modifications included carbamidomethylation of cysteine (+57.021465 Da) and TMT labeling of peptide N-termini and lysine residues (+229.1629 Da). Variable modifications included oxidation of methionine (+15.9949 Da, up to 3 per peptide) and protein N-terminal acetylation (+42.0106 Da, up to 1 per peptide). Peptide length was restricted to 7–50 amino acids, with a peptide mass range of 500–5000 Da. Decoy proteins, common contaminants, immunoglobulins, and keratins were filtered from the final protein list. Protein-level intensities were normalized to the intensity of bait (TTR) in each TMT channel. Quantitation was performed by averaging TMT reporter ion intensities for all spectra assigned to each protein across replicate channels within each treatment condition. Fold change was calculated as the ratio of average bortezomib to average vehicle intensity for each protein. Statistical significance between conditions was assessed by two-tailed Student’s t-test, and p-values were corrected for multiple hypothesis testing using the Benjamini-Hochberg procedure and Storey’s q-value method.^77^ The mass spectrometry proteomics data have been deposited to the ProteomeXchange Consortium via the PRIDE^78^ partner repository with the dataset identifier PXD084196.

## Supporting information

Supplemental Figures and Plasmid Sequences

## DATA AVAILABILITY STATEMENT

The mass spectrometry proteomics data have been deposited to the ProteomeXchange Consortium via the PRIDE partner repository with the dataset identifier PXD084196. The RNA-sequencing data generated in this study have been deposited at Gene Expression Omnibus under accession GSE347460 and are available as of the date of publication. Additional data is provided in Five Supplemental Figures. Complete sequences of unpublished plasmids are provided in Supplemental Information.

## ACKNOWLEDGEMENTS

This work was funded by R01GM134125 (J.G.), R15GM116032 (W.S.), R01GM134327 (P.S.) and the University of California. We are grateful to J. Zhou and the Analytical Chemistry Instrumental Facility for technical assistance with intact protein LC-MS, on instrumentation supported by NSF CHE-1828782. C.C. was supported by the California Louis Stokes Alliance for Minority Participation (NSF#1826900). J.P.C. was supported by a T32 Training grant (T32 ES018827). F.S. was supported by the MARC program (T34 GM149470) and the RISE program and its donors. J.C.Gonzalez was supported by the CMSP program under a grant from the Foundation for California Community Colleges Agreement No. 00005976. mycA/B was a generous gift from the late Y. Kishi (Harvard).

## References

(1) Guo, H.; Xiong, Y.; Witkowski, P.; Cui, J.; Wang, L.; Sun, J.; Lara-Lemus, R.; Haataja, L.; Hutchison, K.; Shan, S.; Arvan, P.; Liu, M. Inefficient Translocation of Preproinsulin Contributes to Pancreatic β Cell Failure and Late-Onset Diabetes. J. Biol. Chem. 2014, 289 (23), 16290–16302. 10.1074/jbc.M114.562355

(2) Lacoste, J.; Haghighi, M.; Haider, S.; Reno, C.; Lin, Z.-Y.; Segal, D.; Qian, W. W.; Xiong, X.; Teelucksingh, T.; Miglietta, E.; Shafqat-Abbasi, H.; Ryder, P. V.; Senft, R.; Cimini, B. A.; Murray, R. R.; Nyirakanani, C.; Hao, T.; McClain, G. G.; Roth, F. P.; Calderwood, M. A.; Hill, D. E.; Vidal, M.; Yi, S. S.; Sahni, N.; Peng, J.; Gingras, A.-C.; Singh, S.; Carpenter, A. E.; Taipale, M. Pervasive Mislocalization of Pathogenic Coding Variants Underlying Human Disorders. Cell 2024, 187 (23), 6725–6741.e13. 10.1016/j.cell.2024.09.003

(3) Hegde, R. S.; Voigt, S.; Lingappa, V. R. Regulation of Protein Topology by Trans-Acting Factors at the Endoplasmic Reticulum. Mol. Cell 1998, 2 (1), 85–91. 10.1016/S1097-2765(00)80116-1

(4) Pleiner, T.; Hazu, M.; Pinton Tomaleri, G.; Nguyen, V. N.; Januszyk, K.; Voorhees, R. M. A Selectivity Filter in the ER Membrane Protein Complex Limits Protein Misinsertion at the ER. J. Cell Biol. 2023, 222 (8), e202212007. 10.1083/jcb.202212007

(5) Fewell, S. W.; Brodsky, J. L. Entry into the Endoplasmic Reticulum: Protein Translocation, Folding and Quality Control. In Madame Curie Bioscience Database [Internet]; Landes Bioscience, 2013.

(6) McKenna, M.; Simmonds, R. E.; High, S. Mechanistic Insights into the Inhibition of Sec61-Dependent Co- and Post-Translational Translocation by Mycolactone. J. Cell Sci. 2016, 129 (7), 1404–1415. 10.1242/jcs.182352

(7) Itskanov, S.; Wang, L.; Junne, T.; Sherriff, R.; Xiao, L.; Blanchard, N.; Shi, W. Q.; Forsyth, C.; Hoepfner, D.; Spiess, M.; Park, E. A Common Mechanism of Sec61 Translocon Inhibition by Small Molecules. Nat. Chem. Biol. 2023, 19 (9), 1063–1071. 10.1038/s41589-023-01337-y

(8) Garrison, J. L.; Kunkel, E. J.; Hegde, R. S.; Taunton, J. A Substrate-Specific Inhibitor of Protein Translocation into the Endoplasmic Reticulum. Nature 2005, 436 (7048), 285–289. 10.1038/nature03821

(9) Demangel, C.; High, S. Sec61 Blockade by Mycolactone: A Central Mechanism in Buruli Ulcer Disease. Biol. Cell 2018, 110 (11), 237–248. 10.1111/boc.201800030

(10) Roboti, P.; O’Keefe, S.; Duah, K. B.; Shi, W. Q.; High, S. Ipomoeassin-F Disrupts Multiple Aspects of Secretory Protein Biogenesis. Sci. Rep. 2021, 11 (1), 11562. 10.1038/s41598-021-91107-4

(11) Kang, S.-W.; Rane, N. S.; Kim, S. J.; Garrison, J. L.; Taunton, J.; Hegde, R. S. Substrate-Specific Translocational Attenuation during ER Stress Defines a Pre-Emptive Quality Control Pathway. Cell 2006, 127 (5), 999–1013. 10.1016/j.cell.2006.10.032

(12) Espinoza, M. F.; Nguyen, K. K.; Sycks, M. M.; Lyu, Z.; Quanrud, G. M.; Montoya, M. R.; Genereux, J. C. Heat Shock Protein Hspa13 Regulates Endoplasmic Reticulum and Cytosolic Proteostasis through Modulation of Protein Translocation. J. Biol. Chem. 2022, 298 (12), 102597. 10.1016/j.jbc.2022.102597

(13) Kadowaki, H.; Nagai, A.; Maruyama, T.; Takami, Y.; Satrimafitrah, P.; Kato, H.; Honda, A.; Hatta, T.; Natsume, T.; Sato, T.; Kai, H.; Ichijo, H.; Nishitoh, H. Pre-Emptive Quality Control Protects the ER from Protein Overload via the Proximity of ERAD Components and SRP. Cell Rep. 2015, 13 (5), 944–956. 10.1016/j.celrep.2015.09.047

(14) Liu, Y.; Gu, Y.; Chen, Y.; Wang, X.; Zhou, G.; Li, J.; Wang, M.; Fang, S.; Yang, Y. Translocational Attenuation Mediated by the PERK-SRP14 Axis Is a Protective Mechanism of Unfolded Protein Response. Cell Rep. 2024, 43 (7), 114402. 10.1016/j.celrep.2024.114402

(15) Hessa, T.; Sharma, A.; Mariappan, M.; Eshleman, H. D.; Gutierrez, E.; Hegde, R. S. Protein Targeting and Degradation Are Coupled for Elimination of Mislocalized Proteins. Nature 2011, 475 (7356), 394–397. 10.1038/nature10181

(16) Rodrigo-Brenni, M. C.; Gutierrez, E.; Hegde, R. S. Cytosolic Quality Control of Mislocalized Proteins Requires RNF126 Recruitment to Bag6. Mol. Cell 2014, 55 (2), 227–237. 10.1016/j.molcel.2014.05.025

(17) Kadowaki, H.; Nishitoh, H. Endoplasmic Reticulum Quality Control by Garbage Disposal. FEBS J. 2019, 286 (2), 232–240. 10.1111/febs.14589

(18) Kasu, Y. A. T.; Arva, A.; Johnson, J.; Sajan, C.; Manzano, J.; Hennes, A.; Haynes, J.; Brower, C. S. BAG6 Prevents the Aggregation of Neurodegeneration-Associated Fragments of TDP43. iScience 2022, 25 (5), 104273. 10.1016/j.isci.2022.104273

(19) Sundaram, A.; Plumb, R.; Appathurai, S.; Mariappan, M. The Sec61 Translocon Limits IRE1α Signaling during the Unfolded Protein Response. eLife 2017, 6, e27187. 10.7554/eLife.27187

(20) Plumb, R.; Zhang, Z.-R.; Appathurai, S.; Mariappan, M. A Functional Link between the Co-Translational Protein Translocation Pathway and the UPR. eLife 2015, 4, e07426. 10.7554/eLife.07426

(21) Zappa, F.; Subramanian, A.; Yang, B.; Conrad, J. E.; Lu, T.-W.; Wang, J.; DeBeaubien, N. A.; Yan, R.; Minopoli, R.; Croll, T.; Tyanova, S.; Costa-Mattioli, M.; Walter, P.; Acosta-Alvear, D. IRE1 Drives a Homeostatic Response to Reduced Protein Influx into the Endoplasmic Reticulum. bioRxiv March 9, 2026, p 2026.03.02.709157. 10.64898/2026.03.02.709157

(22) Morel, J.-D.; Paatero, A. O.; Wei, J.; Yewdell, J. W.; Guenin-Macé, L.; Van Haver, D.; Impens, F.; Pietrosemoli, N.; Paavilainen, V. O.; Demangel, C. Proteomics Reveals Scope of Mycolactone-Mediated Sec61 Blockade and Distinctive Stress Signature*. Mol. Cell. Proteomics 2018, 17 (9), 1750–1765. 10.1074/mcp.RA118.000824

(23) Baron, L.; Paatero, A. O.; Morel, J.-D.; Impens, F.; Guenin-Macé, L.; Saint-Auret, S.; Blanchard, N.; Dillmann, R.; Niang, F.; Pellegrini, S.; Taunton, J.; Paavilainen, V. O.; Demangel, C. Mycolactone Subverts Immunity by Selectively Blocking the Sec61 Translocon. J. Exp. Med. 2016, 213 (13), 2885–2896. 10.1084/jem.20160662

(24) Lyu, Z.; Sycks, M. M.; Espinoza, M. F.; Nguyen, K. K.; Montoya, M. R.; Galapate, C. M.; Mei, L.; Genereux, J. C. Monitoring Protein Import into the Endoplasmic Reticulum in Living Cells with Proximity Labeling. ACS Chem. Biol. 2022, 17 (7), 1963–1977. 10.1021/acschembio.2c00405

(25) Khosravi, A.; Nnamdi, P.; May, A.; Slattery, K.; Sammelson, R. E.; Shi, W. Q. Structural Modifications Reveal Dual Functions of the C-4 Carbonyl Group in the Fatty Acid Chain of Ipomoeassin F. Molecules 2025, 30 (2), 400. 10.3390/molecules30020400

(26) O’Keefe, S.; Bhadra, P.; Duah, K. B.; Zong, G.; Tenay, L.; Andrews, L.; Schneider, H.; Anderson, A.; Hu, Z.; Aljewari, H. S.; Hall, B. S.; Simmonds, R. E.; Helms, V.; High, S.; Shi, W. Q. Synthesis, Biological Evaluation and Docking Studies of Ring-Opened Analogues of Ipomoeassin F. Molecules 2022, 27 (14), 4419. 10.3390/molecules27144419

(27) Song, F.; Fidanze, S.; Benowitz, A. B.; Kishi, Y. Total Synthesis of the Mycolactones. Org. Lett. 2002, 4 (4), 647–650. 10.1021/ol0172828

(28) Kubicek-Sutherland, J. Z.; Swanson, B. I.; Mukundan, H. Conditions for Handling and Optimal Storage of Mycolactone. Methods Mol. Biol. Clifton NJ 2022, 2387, 109–116. 10.1007/978-1-0716-1779-3_12

(29) Förster, B.; Demangel, C.; Thye, T. Mycolactone Induces Cell Death by SETD1B-Dependent Degradation of Glutathione. PLoS Negl. Trop. Dis. 2020. 10.1371/journal.pntd.0008709

(30) Grandjean, J. M. D.; Plate, L.; Morimoto, R. I.; Bollong, M. J.; Powers, E. T.; Wiseman, R. L. Deconvoluting Stress-Responsive Proteostasis Signaling Pathways for Pharmacologic Activation Using Targeted RNA Sequencing. ACS Chem. Biol. 2019, 14 (4), 784–795. 10.1021/acschembio.9b00134

(31) Bush, K. T.; Goldberg, A. L.; Nigam, S. K. Proteasome Inhibition Leads to a Heat-Shock Response, Induction of Endoplasmic Reticulum Chaperones, and Thermotolerance. J. Biol. Chem. 1997, 272 (14), 9086–9092. 10.1074/jbc.272.14.9086

(32) Kim, H.-J.; Joo, H. J.; Kim, Y. H.; Ahn, S.; Chang, J.; Hwang, K.-B.; Lee, D.-H.; Lee, K.-J. Systemic Analysis of Heat Shock Response Induced by Heat Shock and a Proteasome Inhibitor MG132. PLOS ONE 2011, 6 (6), e20252. 10.1371/journal.pone.0020252

(33) Domenger, A.; Choisy, C.; Baron, L.; Mayau, V.; Perthame, E.; Deriano, L.; Arnulf, B.; Bories, J.-C.; Dadaglio, G.; Demangel, C. The Sec61 Translocon Is a Therapeutic Vulnerability in Multiple Myeloma. EMBO Mol. Med. 2022, 14 (3), e14740. 10.15252/emmm.202114740

(34) Hall, B. S.; Hill, K.; McKenna, M.; Ogbechi, J.; High, S.; Willis, A. E.; Simmonds, R. E. The Pathogenic Mechanism of the Mycobacterium Ulcerans Virulence Factor, Mycolactone, Depends on Blockade of Protein Translocation into the ER. PLoS Pathog. 2014, 10 (4), e1004061. 10.1371/journal.ppat.1004061

(35) Hollien, J.; Lin, J. H.; Li, H.; Stevens, N.; Walter, P.; Weissman, J. S. Regulated Ire1-Dependent Decay of Messenger RNAs in Mammalian Cells. J. Cell Biol. 2009, 186 (3), 323–331. 10.1083/jcb.200903014

(36) Ye, J.; Rawson, R. B.; Komuro, R.; Chen, X.; Davé, U. P.; Prywes, R.; Brown, M. S.; Goldstein, J. L. ER Stress Induces Cleavage of Membrane-Bound ATF6 by the Same Proteases That Process SREBPs. Mol. Cell 2000, 6 (6), 1355–1364. 10.1016/S1097-2765(00)00133-7

(37) Sekijima, Y.; Wiseman, R. L.; Matteson, J.; Hammarström, P.; Miller, S. R.; Sawkar, A. R.; Balch, W. E.; Kelly, J. W. The Biological and Chemical Basis for Tissue-Selective Amyloid Disease. Cell 2005, 121 (1), 73–85. 10.1016/j.cell.2005.01.018

(38) Sogawa, A.; Komori, R.; Yanagitani, K.; Ohfurudono, M.; Tsuru, A.; Kadoi, K.; Kimata, Y.; Yoshida, H.; Kohno, K. Signal Sequence-Triage Is Activated by Translocon Obstruction Sensed by an ER Stress Sensor IRE1α. Cell Struct. Funct. 2023, 48 (2), 211–221. 10.1247/csf.23072

(39) Kass, I.; Knaupp, A. S.; Bottomley, S. P.; Buckle, A. M. Conformational Properties of the Disease-Causing Z Variant of Α1-Antitrypsin Revealed by Theory and Experiment. Biophys. J. 2012, 102 (12), 2856–2865. 10.1016/j.bpj.2012.05.023

(40) Christianson, J. C.; Shaler, T. A.; Tyler, R. E.; Kopito, R. R. OS-9 and GRP94 Deliver Mutant Alpha1-Antitrypsin to the Hrd1-SEL1L Ubiquitin Ligase Complex for ERAD. Nat. Cell Biol. 2008, 10 (3), 272–282. 10.1038/ncb1689

(41) Christianson, J. C.; Olzmann, J. A.; Shaler, T. A.; Sowa, M. E.; Bennett, E. J.; Richter, C. M.; Tyler, R. E.; Greenblatt, E. J.; Wade Harper, J.; Kopito, R. R. Defining Human ERAD Networks through an Integrative Mapping Strategy. Nat. Cell Biol. 2012, 14 (1), 93–105. 10.1038/ncb2383

(42) Chen, J. J.; Genereux, J. C.; Qu, S.; Hulleman, J. D.; Shoulders, M. D.; Wiseman, R. L. ATF6 Activation Reduces the Secretion and Extracellular Aggregation of Destabilized Variants of an Amyloidogenic Protein. Chem. Biol. 2014, 21 (11), 1564–1574. 10.1016/j.chembiol.2014.09.009

(43) Legesse, A.; Kaushansky, N.; Braunstein, I.; Saad, H.; Lederkremer, G.; Navon, A.; Stanhill, A. The Role of RNF149 in the Pre-Emptive Quality Control Substrate Ubiquitination. Commun. Biol. 2023, 6 (1), 385. 10.1038/s42003-023-04763-9

(44) Wang, X.; Cattaneo, F.; Ryno, L.; Hulleman, J.; Reixach, N.; Buxbaum, J. N. The Systemic Amyloid Precursor Transthyretin (TTR) Behaves as a Neuronal Stress Protein Regulated by HSF1 in SH-SY5Y Human Neuroblastoma Cells and APP23 Alzheimer’s Disease Model Mice. J. Neurosci. 2014, 34 (21), 7253–7265. 10.1523/JNEUROSCI.4936-13.2014

(45) Lyu, Z.; Genereux, J. Quantitative Measurement of Transthyretin Mistargeting by Proximity Labeling and Parallel Reaction Monitoring. Front. Chem. Biol. 2023, 2. 10.3389/fchbi.2023.1288188

(46) Ciechanover, A.; Schwartz, A. L. The Ubiquitin-Proteasome Pathway: The Complexity and Myriad Functions of Proteins Death. Proc. Natl. Acad. Sci. U. S. A. 1998, 95 (6), 2727–2730.

(47) Krshnan, L.; van de Weijer, M. L.; Carvalho, P. Endoplasmic Reticulum-Associated Protein Degradation. Cold Spring Harb. Perspect. Biol. 2022, 14 (12), a041247. 10.1101/cshperspect.a041247

(48) Hammarström, P.; Wiseman, R. L.; Powers, E. T.; Kelly, J. W. Prevention of Transthyretin Amyloid Disease by Changing Protein Misfolding Energetics. Science 2003, 299 (5607), 713–716. 10.1126/science.1079589

(49) Sifers, R. N.; Brashears-Macatee, S.; Kidd, V. J.; Muensch, H.; Woo, S. L. A Frameshift Mutation Results in a Truncated Alpha 1-Antitrypsin That Is Retained within the Rough Endoplasmic Reticulum. J. Biol. Chem. 1988, 263 (15), 7330–7335. 10.1016/S0021-9258(18)68646-6

(50) Fu, J.; Gao, J.; Liang, Z.; Yang, D. PDI-Regulated Disulfide Bond Formation in Protein Folding and Biomolecular Assembly. Molecules 2021, 26 (1), 171. 10.3390/molecules26010171

(51) Robinson, P. J.; Kanemura, S.; Cao, X.; Bulleid, N. J. Protein Secondary Structure Determines the Temporal Relationship between Folding and Disulfide Formation. bioRxiv March 1, 2019, p 564740. 10.1101/564740

(52) Sun, S.; Li, X.; Mariappan, M. Signal Sequences Encode Information for Protein Folding in the Endoplasmic Reticulum. J. Cell Biol. 2022, 222 (1), e202203070. 10.1083/jcb.202203070

(53) Rutkowski, D. T.; Kang, S.-W.; Goodman, A. G.; Garrison, J. L.; Taunton, J.; Katze, M. G.; Kaufman, R. J.; Hegde, R. S. The Role of p58IPK in Protecting the Stressed Endoplasmic Reticulum. Mol. Biol. Cell 2007, 18 (9), 3681–3691. 10.1091/mbc.e07-03-0272

(54) Sato, T.; Sako, Y.; Sho, M.; Momohara, M.; Suico, M. A.; Shuto, T.; Nishitoh, H.; Okiyoneda, T.; Kokame, K.; Kaneko, M.; Taura, M.; Miyata, M.; Chosa, K.; Koga, T.; Morino-Koga, S.; Wada, I.; Kai, H. STT3B-Dependent Posttranslational N-Glycosylation as a Surveillance System for Secretory Protein. Mol. Cell 2012, 47 (1), 99–110. 10.1016/j.molcel.2012.04.015

(55) Murray, A. N.; Chen, W.; Antonopoulos, A.; Hanson, S. R.; Wiseman, R. L.; Dell, A.; Haslam, S. M.; Powers, D. L.; Powers, E. T.; Kelly, J. W. Enhanced Aromatic Sequons Increase Oligosaccharyltransferase Glycosylation Efficiency and Glycan Homogeneity. Chem. Biol. 2015, 22 (8), 1052–1062. 10.1016/j.chembiol.2015.06.017

(56) Langin, G.; Raffeiner, M.; Biermann, D.; Franz, M.; Spinti, D.; Börnke, F.; Macek, B.; Üstün, S. Proteotoxic Stress Response Is Governed by ER-Associated Sorting of Proteasome Transcriptional Activators. Mol. Cell 2026, 86 (10), 1963–1978.e7. 10.1016/j.molcel.2026.04.004

(57) Yagishita, N.; Aratani, S.; Leach, C.; Amano, T.; Yamano, Y.; Nakatani, K.; Nishioka, K.; Nakajima, T. RING-Finger Type E3 Ubiquitin Ligase Inhibitors as Novel Candidates for the Treatment of Rheumatoid Arthritis. Int. J. Mol. Med. 2012, 30 (6), 1281–1286. 10.3892/ijmm.2012.1129

(58) Sato, T.; Susuki, S.; Suico, M. A.; Miyata, M.; Ando, Y.; Mizuguchi, M.; Takeuchi, M.; Dobashi, M.; Shuto, T.; Kai, H. Endoplasmic Reticulum Quality Control Regulates the Fate of Transthyretin Variants in the Cell. EMBO J. 2007, 26 (10), 2501–2512. 10.1038/sj.emboj.7601685

(59) Bersuker, K.; Brandeis, M.; Kopito, R. R. Protein Misfolding Specifies Recruitment to Cytoplasmic Inclusion Bodies. J. Cell Biol. 2016, 213 (2), 229–241. 10.1083/jcb.201511024

(60) Mauvezin, C.; Neufeld, T. P. Bafilomycin A1 Disrupts Autophagic Flux by Inhibiting Both V-ATPase-Dependent Acidification and Ca-P60A/SERCA-Dependent Autophagosome-Lysosome Fusion. Autophagy 2015, 11 (8), 1437–1438. 10.1080/15548627.2015.1066957

(61) Ronan, B.; Flamand, O.; Vescovi, L.; Dureuil, C.; Durand, L.; Fassy, F.; Bachelot, M.-F.; Lamberton, A.; Mathieu, M.; Bertrand, T.; Marquette, J.-P.; El-Ahmad, Y.; Filoche-Romme, B.; Schio, L.; Garcia-Echeverria, C.; Goulaouic, H.; Pasquier, B. A Highly Potent and Selective Vps34 Inhibitor Alters Vesicle Trafficking and Autophagy. Nat. Chem. Biol. 2014, 10 (12), 1013–1019. 10.1038/nchembio.1681

(62) Pasquier, B. SAR405, a PIK3C3/Vps34 Inhibitor That Prevents Autophagy and Synergizes with MTOR Inhibition in Tumor Cells. Autophagy 2015, 11 (4), 725–726. 10.1080/15548627.2015.1033601

(63) Ward, M. A.; Vangala, J. R.; Kamber Kaya, H. E.; Byers, H. A.; Hosseini, N.; Diaz, A.; Cuervo, A. M.; Kaushik, S.; Radhakrishnan, S. K. Transcription Factor Nrf1 Regulates Proteotoxic Stress-Induced Autophagy. J. Cell Biol. 2024, 223 (6), e202306150. 10.1083/jcb.202306150

(64) Waltho, A.; Popp, O.; Lenz, C.; Pluska, L.; Lambert, M.; Dötsch, V.; Mertins, P.; Sommer, T. K48- and K63-Linked Ubiquitin Chain Interactome Reveals Branch- and Length-Specific Ubiquitin Interactors. Life Sci. Alliance 2024, 7 (8), e202402740. 10.26508/lsa.202402740

(65) Chen, J. J.; Genereux, J. C.; Suh, E. H.; Vartabedian, V. F.; Rius, B.; Qu, S.; Dendle, M. T. A.; Kelly, J. W.; Wiseman, R. L. Endoplasmic Reticulum Proteostasis Influences the Oligomeric State of an Amyloidogenic Protein Secreted from Mammalian Cells. Cell Chem. Biol. 2016, 23 (10), 1282–1293. 10.1016/j.chembiol.2016.09.001

(66) Bulawa, C. E.; Connelly, S.; DeVit, M.; Wang, L.; Weigel, C.; Fleming, J. A.; Packman, J.; Powers, E. T.; Wiseman, R. L.; Foss, T. R.; Wilson, I. A.; Kelly, J. W.; Labaudinière, R. Tafamidis, a Potent and Selective Transthyretin Kinetic Stabilizer That Inhibits the Amyloid Cascade. Proc. Natl. Acad. Sci. 2012, 109 (24), 9629–9634. 10.1073/pnas.1121005109

(67) Ninagawa, S.; Okada, T.; Sumitomo, Y.; Kamiya, Y.; Kato, K.; Horimoto, S.; Ishikawa, T.; Takeda, S.; Sakuma, T.; Yamamoto, T.; Mori, K. EDEM2 Initiates Mammalian Glycoprotein ERAD by Catalyzing the First Mannose Trimming Step. J. Cell Biol. 2014, 206 (3), 347–356. 10.1083/jcb.201404075

(68) Groisman, B.; Shenkman, M.; Ron, E.; Lederkremer, G. Z. Mannose Trimming Is Required for Delivery of a Glycoprotein from EDEM1 to XTP3-B and to Late Endoplasmic Reticulum-Associated Degradation Steps. J. Biol. Chem. 2011, 286 (2), 1292–1300. 10.1074/jbc.M110.154849

(69) Oda, Y.; Okada, T.; Yoshida, H.; Kaufman, R. J.; Nagata, K.; Mori, K. Derlin-2 and Derlin-3 Are Regulated by the Mammalian Unfolded Protein Response and Are Required for ER-Associated Degradation. J. Cell Biol. 2006, 172 (3), 383–393. 10.1083/jcb.200507057

(70) Mishra, S.; Paul, R.; Rani, V.; Ghosh, D. K.; Jain, B. P. Cadmium Toxicity on Endoplasmic Reticulum Functioning. Int. J. Biochem. Mol. Biol. 2024, 15 (4), 107–117. 10.62347/OUDS3732

(71) Woeste, M.; Steller, J.; Hofmann, E.; Kidd, T.; Patel, R.; Connolly, K.; Jayasinghe, M.; Paula, S. Structural Requirements for Inhibitory Effects of Bisphenols on the Activity of the Sarco/Endoplasmic Reticulum Calcium ATPase. Bioorg. Med. Chem. 2013, 21 (13), 3927–3933. 10.1016/j.bmc.2013.04.012

(72) Michelangeli, F.; Ogunbayo, O. A.; Wootton, L. L.; Lai, P. F.; Al-Mousa, F.; Harris, R. M.; Waring, R. H.; Kirk, C. J. Endocrine Disrupting Alkylphenols: Structural Requirements for Their Adverse Effects on Ca2+pumps, Ca2+ Homeostasis & Sertoli TM4 Cell Viability. Chem. Biol. Interact. 2008, 176 (2), 220–226. 10.1016/j.cbi.2008.08.005

(73) He, L.; Diedrich, J.; Chu, Y.-Y.; Yates, J. R., III. Extracting Accurate Precursor Information for Tandem Mass Spectra by RawConverter. Anal. Chem. 2015, 87 (22), 11361–11367. 10.1021/acs.analchem.5b02721

(74) The UniProt Consortium. UniProt: A Worldwide Hub of Protein Knowledge. Nucleic Acids Res. 2019, 47 (D1), D506–D515. 10.1093/nar/gky1049

(75) Peng, Junmin; Elias, Joshua E.; Thoreen, Carson C.; Licklider, Lewis J.; Gygi, Steven P. Evaluation of Multidimensional Chromatography Coupled with Tandem Mass Spectrometry (LC/LC−MS/MS) for Large-Scale Protein Analysis: The Yeast Proteome | Journal of Proteome Research | ACS Publications. J. Proteome Res. 2003. 10.1021/pr025556v

(76) Kong, A. T.; Leprevost, F. V.; Avtonomov, D. M.; Mellacheruvu, D.; Nesvizhskii, A. I. MSFragger: Ultrafast and Comprehensive Peptide Identification in Shotgun Proteomics. Nat. Methods 2017, 14 (5), 513–520. 10.1038/nmeth.4256

(77) Storey, J. D.; Tibshirani, R. Statistical Significance for Genomewide Studies. Proc. Natl. Acad. Sci. 2003, 100 (16), 9440–9445. 10.1073/pnas.1530509100

(78) Perez-Riverol, Y.; Bandla, C.; Kundu, D. J.; Kamatchinathan, S.; Bai, J.; Hewapathirana, S.; John, N. S.; Prakash, A.; Walzer, M.; Wang, S.; Vizcaíno, J. A. The PRIDE Database at 20 Years: 2025 Update. Nucleic Acids Res. 2025, 53 (D1), D543–D553. 10.1093/nar/gkae1011

