## Supplemental Figures and Plasmid Sequences for "N-terminal Trimming Supports Proteostasis Under Secretory Protein Mislocalization Stress"

| **Page** | **Contents** |
| --- | --- |
| **S-2** | **Table of Contents** |
| **S-3** | **Supplemental Figure S1** |
| **S-5** | **Supplemental Figure S2** |
| **S-6** | **Supplemental Figure S3** |
| **S-8** | **Supplemental Figure S4** |
| **S-10** | **Supplemental Figure S5** |
| **S-13** | **Plasmid Sequences** |

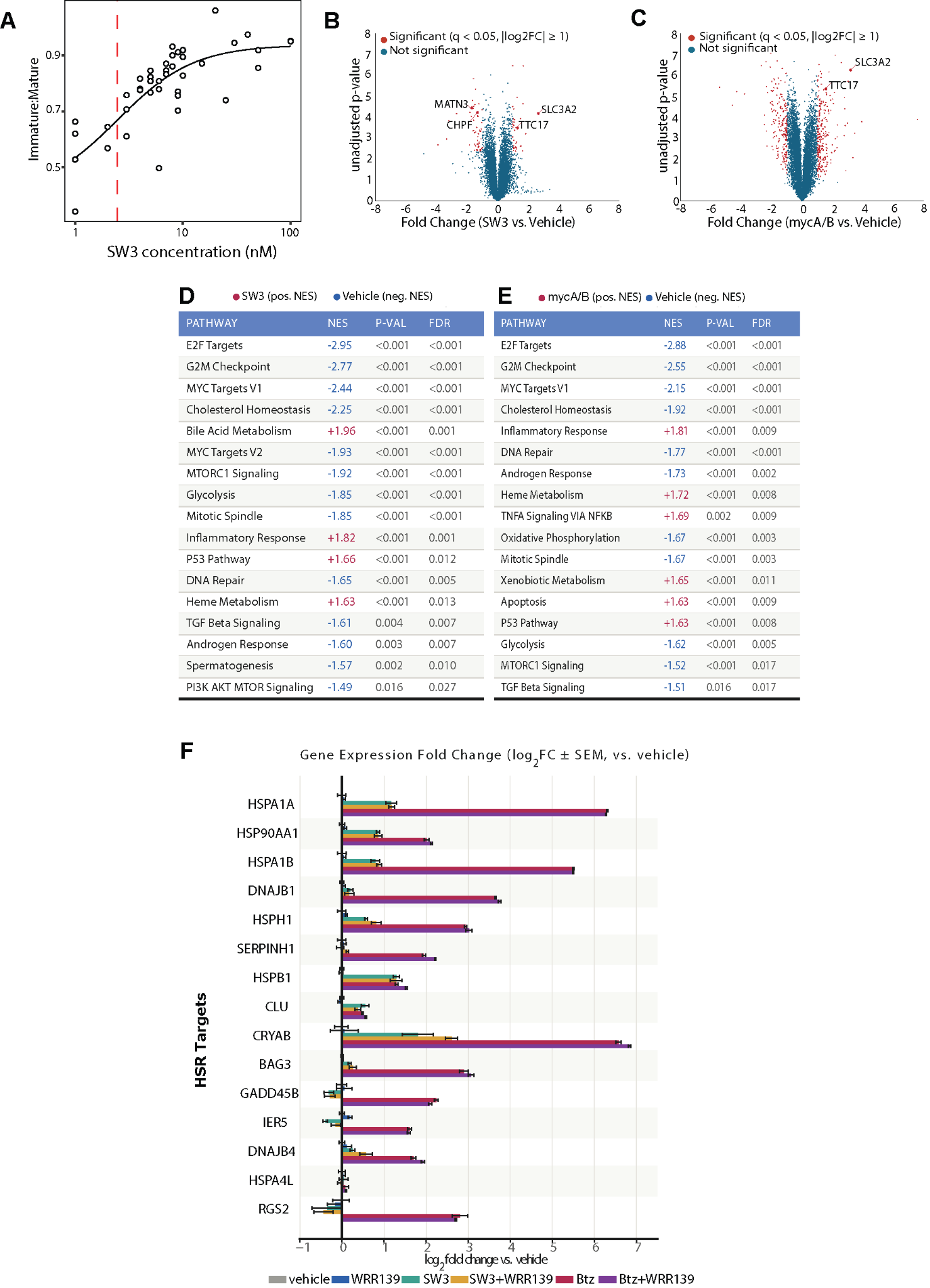

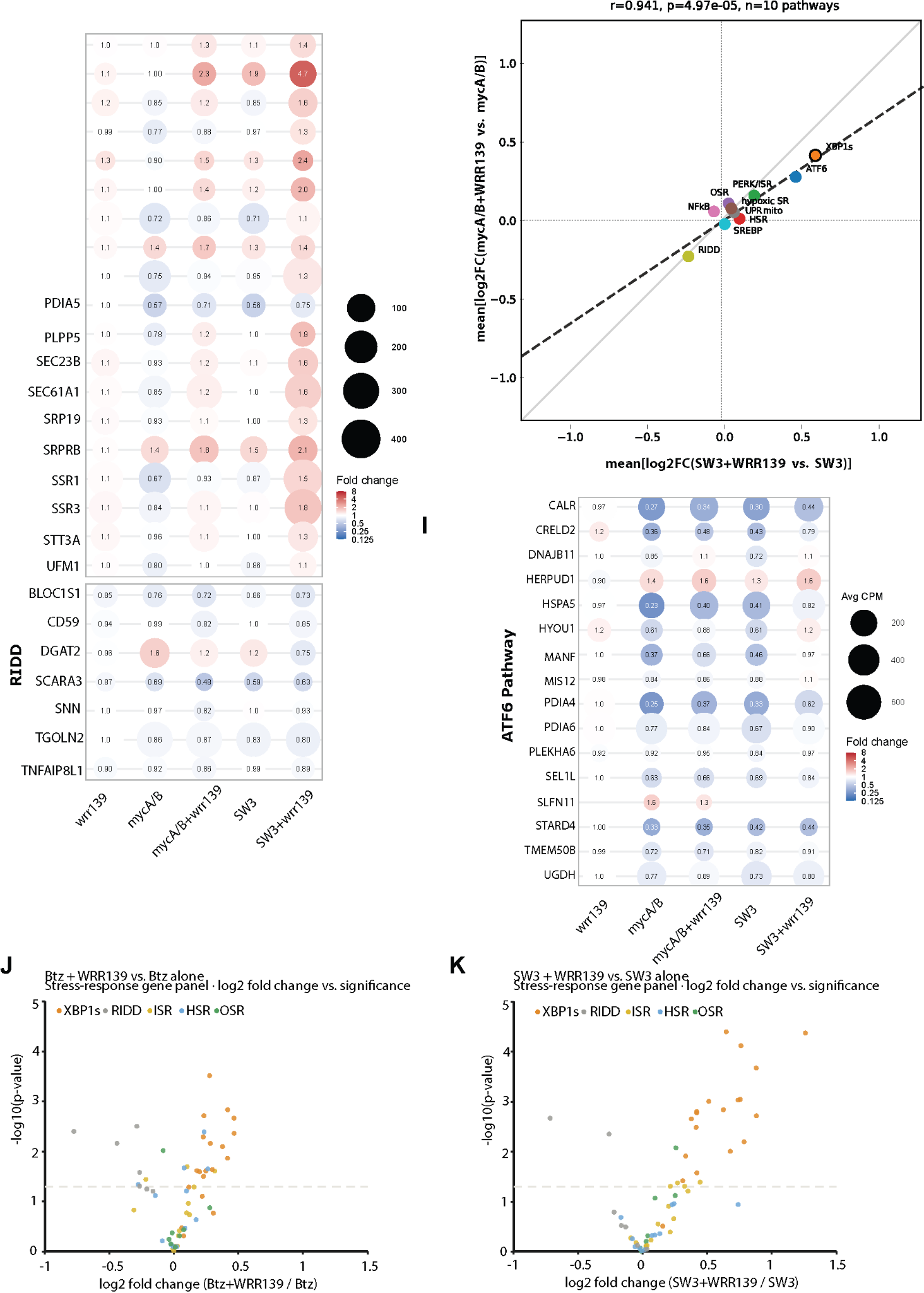

**Figure S1. A.** Dose response of Ipomoeassin F analog SW3–induced TTR mislocalization. HEK293T cells expressing ^Flag^TTR were treated with increasing concentrations of SW3 or mycA/B for 16 h. Quantification of immature (based on signal sequence cleavage)/mature TTR ratio vs inhibitor concentration used to determine EC₅₀. **B.** Volcano plot showing differential gene expression across 12,175 detected protein-coding genes in HepG2 cells treated with SW3 (50 nM, 16 h) versus vehicle (n = 3). Significance was determined using Storey’s modification of the method of Benjamini-Hochberg. **C.** Volcano plot showing differential gene expression across 16,826 detected protein-coding genes in HepG2 cells treated with mycA/B (25 nM, 16 h) versus vehicle (n = 3).Significance in both panels was determined using Storey’s modification of the method of Benjamini-Hochberg. **D.** Gene set enrichment analysis (GSEA) of stress pathways comparing SW3 (50nM) treatment to vehicle. **E.** GSEA of stress pathways comparing mycA/B-treated vs. vehicle-treated HepG2 cells. Normalized Enrichment Score (NES) reflects the degree to which a pathway's genes are shifted toward one condition relative to random chance, correcting for gene set size; positive values indicate enrichment in treatment, negative values indicate enrichment in vehicle. Pathways shown passed a significance threshold of FDR q < 0.05. **F.** Relative transcript levels for target genes as determined by RNAseq for HSR targets genes in HEPG2 cells treated for 16 h with 50 nM SW3, 1 µM Btz, 10 µM WRR139, or vehicle as indicated. Bars represent log₂ fold change ± SEM relative to vehicle (n = 3 biological replicates per condition). **G.** Fold change back-calculated from log_2_FC in mRNA expression relative to each dataset's own vehicle control as determined by RNAseq for IRE1 (RIDD, XBP1) targets genes in HepG2 cells treated for 16 h with 50 nM SW3, 25 nM mycA/B, 10 µM WRR139, or vehicle as indicated. **H.** Relative transcript levels for target genes as determined by RNAseq for ATF6 targets genes in HEPG2 cells treated for 16 h with 20 nM SW3, 1 µM Btz, 10 µM WRR139, or vehicle as indicated. **I.** Gene-level changes induced by WRR139 are plotted as mean[log_2_FC(SW3+WRR139 vs. SW3)] versus mean[log_2_FC(SW3+WRR139 vs. SW3)]. Each point represents a gene, colored by pathway. The dotted line indicates equal WRR139 effect across conditions, and the dashed line shows the linear regression fit. Pearson correlation (*r*) and associated p-value are indicated. **J-K.** Volcano plots illustrating how expression of genes associated with cellular stress responses changes with WRR139 treatment against a background of vehicle, Btz, or SW3 treatment.

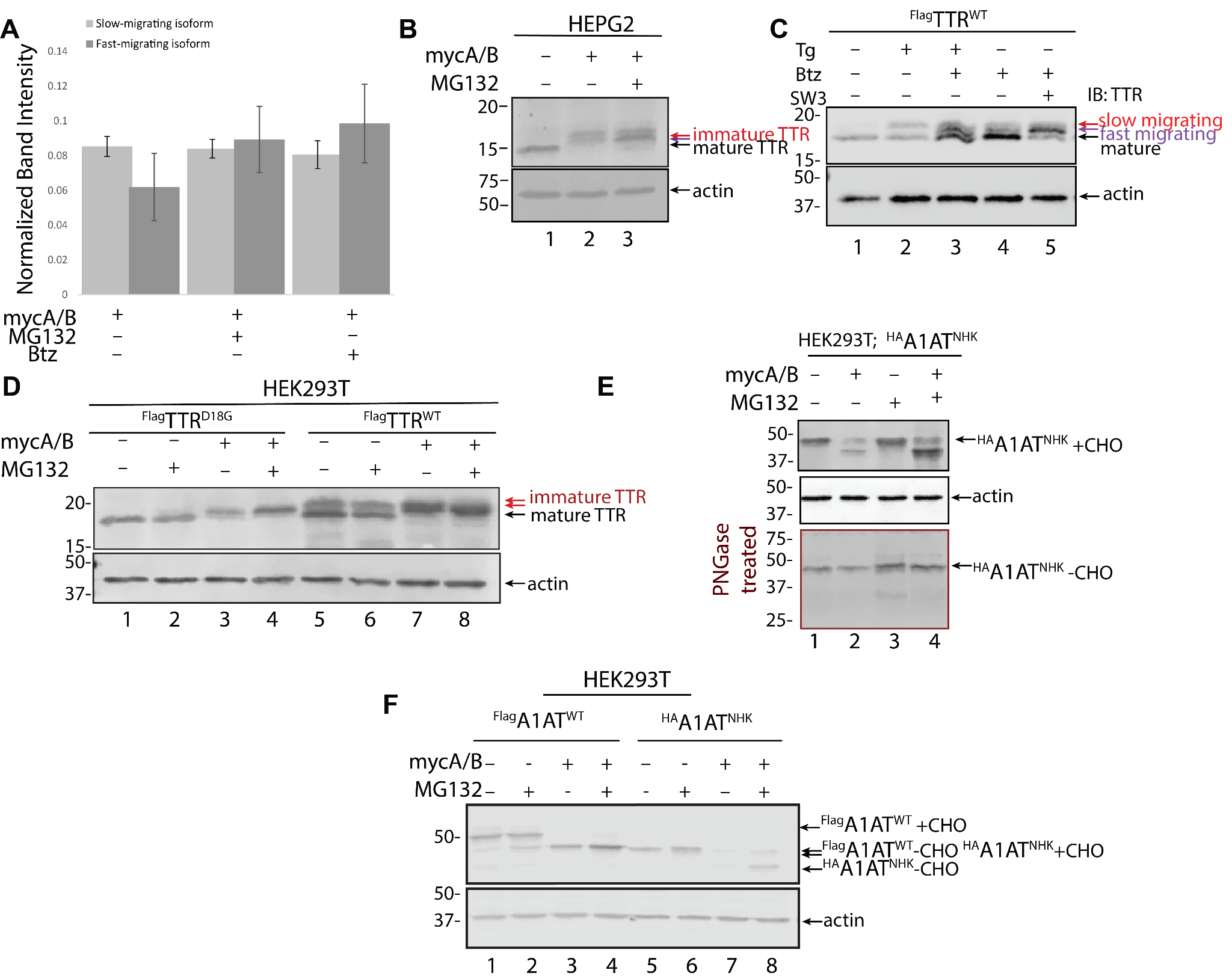

**Figure S2: Western Blot analysis of lysates from HEK293T Cells overexpressing TTR.** **A.** Quantification of immature TTR isoforms under Sec61 inhibition and two different proteasome inhibitors (MG132 or Btz; 1 µM 16 h). Densitometric analysis of the immature TTR isoforms across inhibitor treatments. **B.** Endogenous-context validation in HEPG2 cells. HEPG2 cells (expressing endogenous, untagged TTR) were treated with mycA/B ± MG132 for 16 h, error bars represent SD, *n* = 4. Representative immunoblot shows two immature TTR isoforms similar to those observed in overexpression systems, confirming tag-independent behavior. **C.** Representative immunoblot of SDS-PAGE-separated lysates from HEK239T cells overexpressing ^FLAG^TTR and treated for 16 h with 50 nM Tg, 1 µM Btz, and/or 10 nM SW3 as indicated. **D.** HEK293T cells expressing ^Flag^TTR^WT^ or ^Flag^TTR^D18G^ were treated with mycA/B ± MG132 for 16 h. The fast-migrating (intact) isoform is strongly restored by proteasome inhibition; the slow-migrating N-terminally trimmed isoform remains largely persistent. **E.** Immunoblot of ^Flag^A1AT^WT^ and ^Flag^A1AT^NHK^ species in HEK293T cells under the indicated treatments. The bottom panel of the immunoblot is treated with PNGase (indicated in red border). *n* = 2. **F.** HEK293T cells expressing ^Flag^A1AT^WT^ or the destabilized patient variant ^Flag^A1AT^NHK^ were treated with mycA/B ± MG132 for 16 h, *n* = 4. MycA/B eliminates ER glycosylation (^Flag^A1AT^WT^+CHO, carbohydrate), shifting A1AT to lower molecular-weight species (^Flag^A1AT^WT^-CHO). ^Flag^A1AT^NHK^ is strongly degraded upon mislocalization, and proteasome inhibition restores its cytosolic (^Flag^A1AT^NHK^ -CHO) immature band.

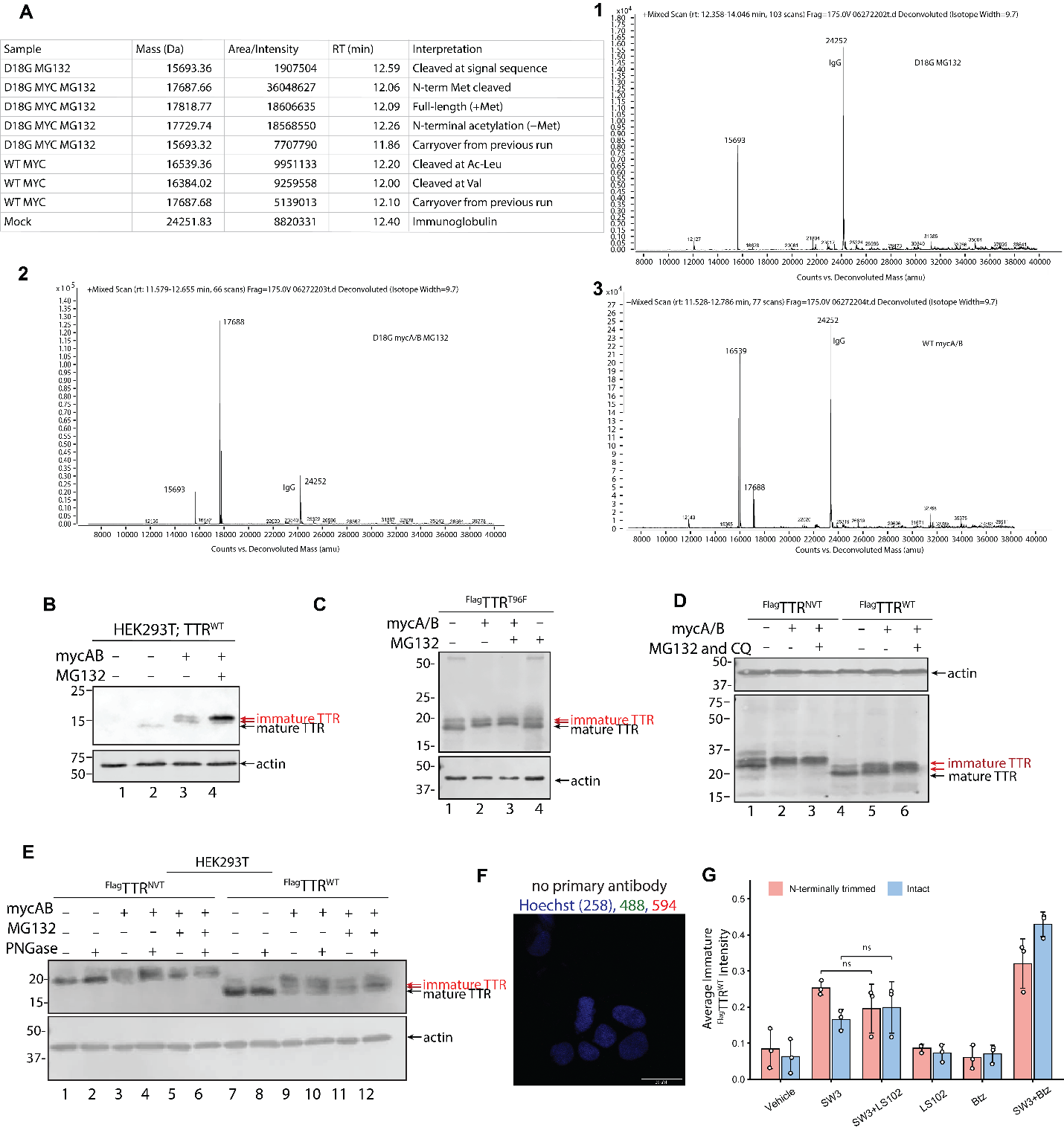

**Figure S3. Validation of N-terminal trimming and glycosylation reporters under Sec61 inhibition. A.** Observed deconvoluted masses, relative intensities, retention times, and assignments for **(1)** mature, **(2)** intact intermediate, **(3)** N-terminally trimmed species are shown. TTR isoforms isolated from HEK293T cells expressing ^Flag^TTR variants under mycA/B (25 nM, 16 h) and MG132 (1 µM, 16 h) treatment. **B.** Representative western blot of HEK293T overexpressing non-FLAG-tagged TTR treated as indicated, *n* = 3. **C.** Representative western blot of HEK293T cells expressing ^Flag^TTR^T96F^ treated with mycA/B and MG132, *n* = 3. **D.** Representative western blot of HEK293T cells expressing ^Flag^TTR^NVT^. The C-terminal glycosylation reporter (GS linker + NVT sequon: GGCAGCGGCAGCGGTAGTGGTAGTGGCAGCAACGTGACCAGCAGCTAGTAA, *n* = 3). **E.** Same samples as in (D) analyzed with or without PNGase F treatment. **F.** Representative immunofluorescence micrograph of HEK293T cells incubated with secondary antibody alone showing minimal background and nuclei stained with Hoechst. **G.** Quantification of immunoblots from HEK293T cell lysates expressing ^FLAG^TTR^WT^ and treated with 10 µM LS102, 1 µM Btz, or 15 nM as indicated. Error bars represent S.D. and normalized to ponceau total lysate blot.

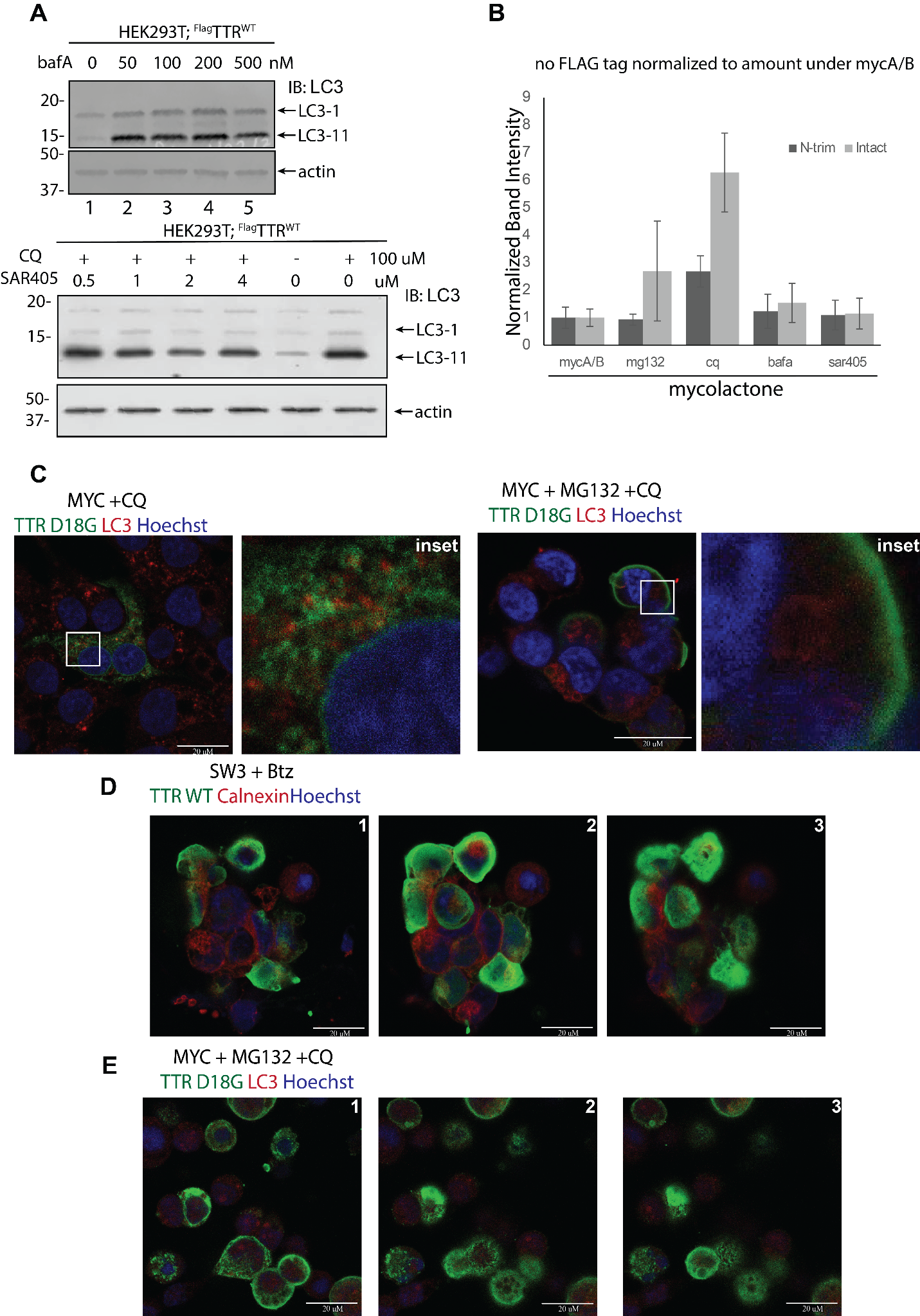

**Figure S4A**. Representative immunoblots of lysates from HEK293T treated with varying concentrations of the autophagy inhibitors as indicated. **B.** Quantification of mislocalized non-Flag-TTR isoforms in HEK293T cells treated for 16 h with mycolactone (25 nM), MG132 (1 µM), chloroquine (100 µM), bafilomycin A1 (100 nM), or SAR405 (1 µM). Band intensities for the N-terminally trimmed and intact intermediate TTR isoforms were normalized to the mycolactone condition. Data represent mean for *n* = 3 biological replicates. **C.** Cells were stained with anti-TTR (magenta), anti-LAMP1 (green), and Hoechst (blue). Scale bar: 20 µm. **D.** Confocal z-stacks of WT TTR under combined Sec61 inhibition and proteasome inhibition. Bottom of cell on the left (panel 1) and top of cell to the right (panel 3). Scale bar: 20 µm. **E.** Confocal z-stacks of D18G TTR under combined Sec61 inhibition, proteasome inhibition, and CQ treatment. Bottom of cell on the left (panel 1) and top of cell to the right (panel 3). Scale bar: 20 µm.

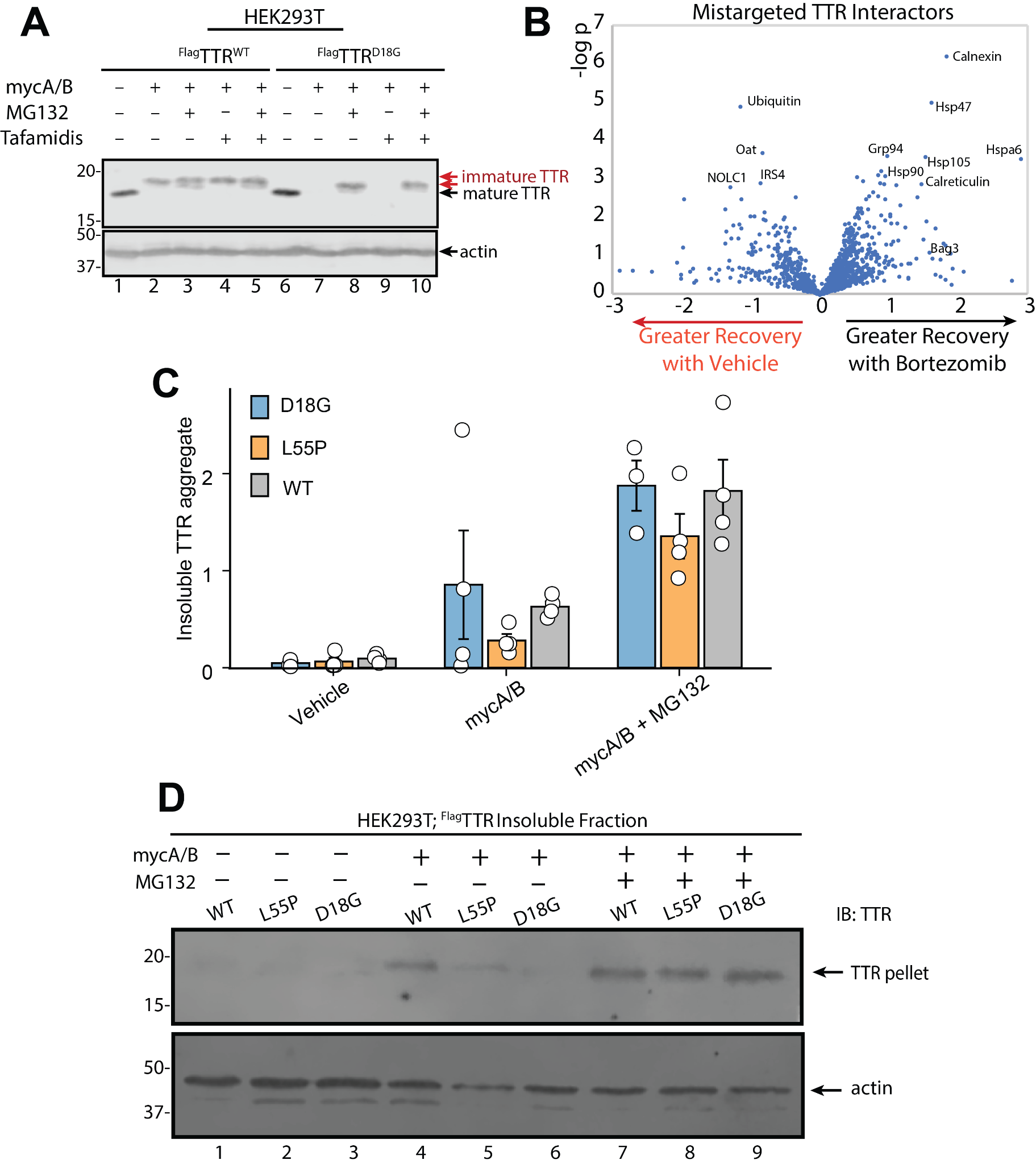

**Figure S5: A. Western Blot analysis of lysates from HEK293T Cells overexpressing TTR.** Cells were treated for 16 h with 25 nM mycAB; 1µM MG132; 10 µM Tafadimis. Antibody for TTR is shown on 800 channel only. **B.** Volcano plot of TTR-associated proteins identified by IP-MS (TMT-based quantification) from ^FLAG^TTR^WT^ immunoprecipitates ± bortezomib (Btz) treatment, following DSP crosslinking; abundance normalized to immunoprecipitated TTR. X-axis: log2 fold change in recovery (Btz vs. vehicle). Y-axis: −log10(p). Btz selectively stabilizes intact TTR without affecting the N-terminally trimmed population, proteins enriched with Btz treatment (right) represent preferential intact-TTR interactors. Selected chaperones are labeled (e.g., calnexin, calreticulin, Hsp90, Hsp105, Grp94, Hspa6, Bag3). **C.** Quantification of insoluble TTR aggregates from panel D. Value 6.95 from D18G mycA/B + MG132 group was identified as a statistical outlier by Grubbs’ test (p = 0.0205) and was excluded. Normalization was done to the total lysate under mycA/B + Btz condition to account for all sample inputted in the aggregation assay. Bars represent the mean; dots indicate individual biological replicates. Bars represent mean ± SEM; circles represent individual biological replicates (n = 4). **D.** Immunoblot of insoluble pellets of TTR variants (WT, L55P, D18G) under mislocalization.

**Signal sequence_FT_TTR_WT**

Annotation Key :

FLAG tag

Human A1AT WT

CMV promoter

CMV enhancer

GGCGTAATCTGCTGCTTGCAAACAAAAAAACCACCGCTACCAGCGGTGGTTTGTTTGCCGGATCAAGAGCTACCAACTCTTTTTCCGAAGGTAACTGGCTTCAGCAGAGCGCAGATACCAAATACTGTCCTTCTAGTGTAGCCGTAGTTAGGCCACCACTTCAAGAACTCTGTAGCACCGCCTACATACCTCGCTCTGCTAATCCTGTTACCAGTGGCTGCTGCCAGTGGCGATAAGTCGTGTCTTACCGGGTTGGACTCAAGACGATAGTTACCGGATAAGGCGCAGCGGTCGGGCTGAACGGGGGGTTCGTGCACACAGCCCAGCTTGGAGCGAACGACCTACACCGAACTGAGATACCTACAGCGTGAGCATTGAGAAAGCGCCACGCTTCCCGAAGGGAGAAAGGCGGACAGGTATCCGGTAAGCGGCAGGGTCGGAACAGGAGAGCGCACGAGGGAGCTTCCAGGGGGAAACGCCTGGTATCTTTATAGTCCTGTCGGGTTTCGCCACCTCTGACTTGAGCGTCGATTTTTGTGATGCTCGTCAGGGGGGCGGAGCCTATGGAAAAACGCCAGCAACGCAAGCTAGCTTCTAGCTAGAAATTGTAAACGTTAATATTTTGTTAAAATTCGCGTTAAATTTTTGTTAAATCAGCTCATTTTTTAACCAATAGGCCGAAATCGGCAAAATCCCTTATAAATCAAAAGAATAGCCCGAGATAGGGTTGAGTGTTGTTCCAGTTTGGAACAAGAGTCCACTATTAAAGAACGTGGACTCCAACGTCAAAGGGCGAAAAACCGTCTATCAGGGCGATGGCCGCCCACTACGTGAACCATCACCCAAATCAAGTTTTTTGGGGTCGAGGTGCCGTAAAGCACTAAATCGGAACCCTAAAGGGAGCCCCCGATTTAGAGCTTGACGGGGAAAGCCGGCGAACGTGGCGAGAAAGGAAGGGAAGAAAGCGAAAGGAGCGGGCGCTAGGGCGCTGGCAAGTGTAGCGGTCACGCTGCGCGTAACCACCACACCCGCCGCGCTTAATGCGCCGCTACAGGGCGCGTACTATGGTTGCTTTGACGAGACCGTATAACGTGCTTTCCTCGTTGGAATCAGAGCGGGAGCTAAACAGGAGGCCGATTAAAGGGATTTTAGACAGGAACGGTACGCCAGCTGGATTACCGCGGTCTTTCTCAACGTAACACTTTACAGCGGCGCGTCATTTGATATGATGCGCCCCGCTTCCCGATAAGGGAGCAGGCCAGTAAAAGCATTACCCGTGGTGGGGTTCCCGAGCGGCCAAAGGGAGCAGACTCTAAATCTGCCGTCATCGACTTCGAAGGTTCGAATCCTTCCCCCACCACCATCACTTTCAAAAGTCCGAAAGAATCTGCTCCCTGCTTGTGTGTTGGAGGTCGCTGAGTAGTGCGCGAGTAAAATTTAAGCTACAACAAGGCAAGGCTTGACCGACAATTGCATGAAGAATCTGCTTAGGGTTAGGCGTTTTGCGCTGCTTCGCGATGTACGGGCCAGATATACGCGTTGACATTGATTATTGACTAGTTATTAATAGTAATCAATTACGGGGTCATTAGTTCATAGCCCATATATGGAGTTCCGCGTTACATAACTTACGGTAAATGGCCCGCCTGGCTGACCGCCCAACGACCCCCGCCCATTGACGTCAATAATGACGTATGTTCCCATAGTAACGCCAATAGGGACTTTCCATTGACGTCAATGGGTGGACTATTTACGGTAAACTGCCCACTTGGCAGTACATCAAGTGTATCATATGCCAAGTACGCCCCCTATTGACGTCAATGACGGTAAATGGCCCGCCTGGCATTATGCCCAGTACATGACCTTATGGGACTTTCCTACTTGGCAGTACATCTACGTATTAGTCATCGCTATTACCATGGTGATGCGGTTTTGGCAGTACATCAATGGGCGTGGATAGCGGTTTGACTCACGGGGATTTCCAAGTCTCCACCCCATTGACGTCAATGGGAGTTTGTTTTGGCACCAAAATCAACGGGACTTTCCAAAATGTCGTAACAACTCCGCCCCATTGACGCAAATGGGCGGTAGGCGTGTACGGTGGGAGGTCTATATAAGCAGAGCTCTCTGGCTAACTAGAGAACCCACTGCTTACTGGCTTATCGAAATTAATACGACTCACTATAGGGAGACCCAAGCTTGGTACCGAGCTCGGATCCGCCATGGCTTCTCATCGTCTGCTCCTCCTCTGCCTTGCTGGACTGGTATTTGTGTCTGAGGCTGACTACAAAGACGATGACGATAAGGATTATAAGGATGACGATGACAAGGGTCCTACGGGCACCGGTGAATCCAAGTGTCCTCTGATGGTCAAAGTTCTAGATGCTGTCCGAGGCAGTCCTGCCATCAATGTGGCCGTGCATGTGTTCAGAAAGGCTGCTGATGACACCTGGGAGCCATTTGCCTCTGGGAAAACCAGTGAGTCTGGAGAGCTGCATGGGCTCACAACTGAGGAGGAATTTGTAGAAGGGATATACAAAGTGGAAATAGACACCAAATCTTACTGGAAGGCACTTGGCATCTCCCCATTCCATGAGCATGCAGAGGTGGTATTCACAGCCAACGACTCCGGCCCCCGCCGCTACACCATTGCCGCCCTGCTGAGCCCCTACTCCTATTCCACCACGGCTGTCGTCACCAATCCCAAGGAATGAATTCTGCAGATATCCATCACACTGGCGGCCGCTCGAGCATGCATCTAGAGGGCCCTATTCTATAGTGTCACCTAAATGCTAGAGGATCTTTGTGAAGGAACCTTACTTCTGTGGTGTGACATAATTGGACAAACTACCTACAGAGATTTAAAGCTCTAAGGTAAATATAAAATTTTTAAGTGTATAATGTGTTAAACTACTGATTCTAATTGTTTGTGTATTTTAGATTCCAACCTATGGAACTGATGAATGGGAGCAGTGGTGGAATGCCTTTAATGAGGAAAACCTGTTTTGCTCAGAAGAAATGCCATCTAGTGATGATGAGGCTACTGCTGACTCTCAACATTCTACTCCTCCAAAAAAGAAGAGAAAGGTAGAAGACCCCAAGGACTTTCCTTCAGAATTGCTAAGTTTTTTGAGTCATGCTGTGTTTAGTAATAGAACTCTTGCTTGCTTTGCTATTTACACCACAAAGGAAAAAGCTGCACTGCTATACAAGAAAATTATGGAAAAATATTTGATGTATAGTGCCTTGACTAGAGATCATAATCAGCCATACCACATTTGTAGAGGTTTTACTTGCTTTAAAAAACCTCCCACACCTCCCCCTGAACCTGAAACATAAAATGAATGCAATTGTTGTTGTTAACTTGTTTATTGCAGCTTATAATGGTTACAAATAAAGCAATAGCATCACAAATTTCACAAATAAAGCATTTTTTTCACTGCATTCTAGTTGTGGTTTGTCCAAACTCATCAATGTATCTTATCATGTCTGGATCATCCCGCCATGGTATCAACGCCATATTTCTATTTACAGTAGGGACCTCTTCGTTGTGTAGGTACCGCTGTATTCCTAGGGAAATAGTAGAGGCACCTTGAACTGTCTGCATCAGCCATATAGCCCCCGCTGTTCGACTTACAAACACAGGCACAGTACTGACAAACCCATACACCTCCTCTGAAATACCCATAGTTGCTAGGGCTGTCTCCGAACTCATTACACCCTCCAAAGTCAGAGCTGTAATTTCGCCATCAAGGGCAGCGAGGGCTTCTCCAGATAAAATAGCTTCTGCCGAGAGTCCCGTAAGGGTAGACACTTCAGCTAATCCCTCGATGAGGTCTACTAGAATAGTCAGTGCGGCTCCCATTTTGAAAATTCACTTACTTGATCAGCTTCAGAAGATGGCGGAGGGCCTCCAACACAGTAATTTTCCTCCCGACTCTTAAAATAGAAAATGTCAAGTCAGTTAAGCAGGAAGTGGACTAACTGACGCAGCTGGCCGTGCGACATCCTCTTTTAATTAGTTGCTAGGCAACGCCCTCCAGAGGGCGTGTGGTTTTGCAAGAGGAAGCAAAAGCCTCTCCACCCAGGCCTAGAATGTTTCCACCCAATCATTACTATGACAACAGCTGTTTTTTTTAGTATTAAGCAGAGGCCGGGGACCCCTGGGCCCGCTTACTCTGGAGAAAAAGAAGAGAGGCATTGTAGAGGCTTCCAGAGGCAACTTGTCAAAACAGGACTGCTTCTATTTCTGTCACACTGTCTGGCCCTGTCACAAGGTCCAGCACCTCCATACCCCCTTTAATAAGCAGTTTGGGAACGGGTGCGGGTCTTACTCCGCCCATCCCGCCCCTAACTCCGCCCAGTTCCGCCCATTCTCCGCCCCATGGCTGACTAATTTTTTTTATTTATGCAGAGGCCGAGGCCGCCTCGGCCTCTGAGCTATTCCAGAAGTAGTGAGGAGGCTTTTTTGGAGGCCTAGGCTTTTGCAAAAAGCTAATTC

**Signal sequence_FT_TTR_D18G**

Annotation Key : 5’->3’

FLAG tag

Human A1AT WT

CMV promoter

CMV enhancer

GGCGTAATCTGCTGCTTGCAAACAAAAAAACCACCGCTACCAGCGGTGGTTTGTTTGCCGGATCAAGAGCTACCAACTCTTTTTCCGAAGGTAACTGGCTTCAGCAGAGCGCAGATACCAAATACTGTCCTTCTAGTGTAGCCGTAGTTAGGCCACCACTTCAAGAACTCTGTAGCACCGCCTACATACCTCGCTCTGCTAATCCTGTTACCAGTGGCTGCTGCCAGTGGCGATAAGTCGTGTCTTACCGGGTTGGACTCAAGACGATAGTTACCGGATAAGGCGCAGCGGTCGGGCTGAACGGGGGGTTCGTGCACACAGCCCAGCTTGGAGCGAACGACCTACACCGAACTGAGATACCTACAGCGTGAGCATTGAGAAAGCGCCACGCTTCCCGAAGGGAGAAAGGCGGACAGGTATCCGGTAAGCGGCAGGGTCGGAACAGGAGAGCGCACGAGGGAGCTTCCAGGGGGAAACGCCTGGTATCTTTATAGTCCTGTCGGGTTTCGCCACCTCTGACTTGAGCGTCGATTTTTGTGATGCTCGTCAGGGGGGCGGAGCCTATGGAAAAACGCCAGCAACGCAAGCTAGCTTCTAGCTAGAAATTGTAAACGTTAATATTTTGTTAAAATTCGCGTTAAATTTTTGTTAAATCAGCTCATTTTTTAACCAATAGGCCGAAATCGGCAAAATCCCTTATAAATCAAAAGAATAGCCCGAGATAGGGTTGAGTGTTGTTCCAGTTTGGAACAAGAGTCCACTATTAAAGAACGTGGACTCCAACGTCAAAGGGCGAAAAACCGTCTATCAGGGCGATGGCCGCCCACTACGTGAACCATCACCCAAATCAAGTTTTTTGGGGTCGAGGTGCCGTAAAGCACTAAATCGGAACCCTAAAGGGAGCCCCCGATTTAGAGCTTGACGGGGAAAGCCGGCGAACGTGGCGAGAAAGGAAGGGAAGAAAGCGAAAGGAGCGGGCGCTAGGGCGCTGGCAAGTGTAGCGGTCACGCTGCGCGTAACCACCACACCCGCCGCGCTTAATGCGCCGCTACAGGGCGCGTACTATGGTTGCTTTGACGAGACCGTATAACGTGCTTTCCTCGTTGGAATCAGAGCGGGAGCTAAACAGGAGGCCGATTAAAGGGATTTTAGACAGGAACGGTACGCCAGCTGGATTACCGCGGTCTTTCTCAACGTAACACTTTACAGCGGCGCGTCATTTGATATGATGCGCCCCGCTTCCCGATAAGGGAGCAGGCCAGTAAAAGCATTACCCGTGGTGGGGTTCCCGAGCGGCCAAAGGGAGCAGACTCTAAATCTGCCGTCATCGACTTCGAAGGTTCGAATCCTTCCCCCACCACCATCACTTTCAAAAGTCCGAAAGAATCTGCTCCCTGCTTGTGTGTTGGAGGTCGCTGAGTAGTGCGCGAGTAAAATTTAAGCTACAACAAGGCAAGGCTTGACCGACAATTGCATGAAGAATCTGCTTAGGGTTAGGCGTTTTGCGCTGCTTCGCGATGTACGGGCCAGATATACGCGTTGACATTGATTATTGACTAGTTATTAATAGTAATCAATTACGGGGTCATTAGTTCATAGCCCATATATGGAGTTCCGCGTTACATAACTTACGGTAAATGGCCCGCCTGGCTGACCGCCCAACGACCCCCGCCCATTGACGTCAATAATGACGTATGTTCCCATAGTAACGCCAATAGGGACTTTCCATTGACGTCAATGGGTGGACTATTTACGGTAAACTGCCCACTTGGCAGTACATCAAGTGTATCATATGCCAAGTACGCCCCCTATTGACGTCAATGACGGTAAATGGCCCGCCTGGCATTATGCCCAGTACATGACCTTATGGGACTTTCCTACTTGGCAGTACATCTACGTATTAGTCATCGCTATTACCATGGTGATGCGGTTTTGGCAGTACATCAATGGGCGTGGATAGCGGTTTGACTCACGGGGATTTCCAAGTCTCCACCCCATTGACGTCAATGGGAGTTTGTTTTGGCACCAAAATCAACGGGACTTTCCAAAATGTCGTAACAACTCCGCCCCATTGACGCAAATGGGCGGTAGGCGTGTACGGTGGGAGGTCTATATAAGCAGAGCTCTCTGGCTAACTAGAGAACCCACTGCTTACTGGCTTATCGAAATTAATACGACTCACTATAGGGAGACCCAAGCTTGGTACCGAGCTCGGATCCGCCATGGCTTCTCATCGTCTGCTCCTCCTCTGCCTTGCTGGACTGGTATTTGTGTCTGAGGCTGACTACAAAGACGATGACGATAAGGATTATAAGGATGACGATGACAAGGGTCCTACGGGCACCGGTGAATCCAAGTGTCCTCTGATGGTCAAAGTTCTAGGTGCTGTCCGAGGCAGTCCTGCCATCAATGTGGCCGTGCATGTGTTCAGAAAGGCTGCTGATGACACCTGGGAGCCATTTGCCTCTGGGAAAACCAGTGAGTCTGGAGAGCTGCATGGGCTCACAACTGAGGAGGAATTTGTAGAAGGGATATACAAAGTGGAAATAGACACCAAATCTTACTGGAAGGCACTTGGCATCTCCCCATTCCATGAGCATGCAGAGGTGGTATTCACAGCCAACGACTCCGGCCCCCGCCGCTACACCATTGCCGCCCTGCTGAGCCCCTACTCCTATTCCACCACGGCTGTCGTCACCAATCCCAAGGAATGAATTCTGCAGATATCCATCACACTGGCGGCCGCTCGAGCATGCATCTAGAGGGCCCTATTCTATAGTGTCACCTAAATGCTAGAGGATCTTTGTGAAGGAACCTTACTTCTGTGGTGTGACATAATTGGACAAACTACCTACAGAGATTTAAAGCTCTAAGGTAAATATAAAATTTTTAAGTGTATAATGTGTTAAACTACTGATTCTAATTGTTTGTGTATTTTAGATTCCAACCTATGGAACTGATGAATGGGAGCAGTGGTGGAATGCCTTTAATGAGGAAAACCTGTTTTGCTCAGAAGAAATGCCATCTAGTGATGATGAGGCTACTGCTGACTCTCAACATTCTACTCCTCCAAAAAAGAAGAGAAAGGTAGAAGACCCCAAGGACTTTCCTTCAGAATTGCTAAGTTTTTTGAGTCATGCTGTGTTTAGTAATAGAACTCTTGCTTGCTTTGCTATTTACACCACAAAGGAAAAAGCTGCACTGCTATACAAGAAAATTATGGAAAAATATTTGATGTATAGTGCCTTGACTAGAGATCATAATCAGCCATACCACATTTGTAGAGGTTTTACTTGCTTTAAAAAACCTCCCACACCTCCCCCTGAACCTGAAACATAAAATGAATGCAATTGTTGTTGTTAACTTGTTTATTGCAGCTTATAATGGTTACAAATAAAGCAATAGCATCACAAATTTCACAAATAAAGCATTTTTTTCACTGCATTCTAGTTGTGGTTTGTCCAAACTCATCAATGTATCTTATCATGTCTGGATCATCCCGCCATGGTATCAACGCCATATTTCTATTTACAGTAGGGACCTCTTCGTTGTGTAGGTACCGCTGTATTCCTAGGGAAATAGTAGAGGCACCTTGAACTGTCTGCATCAGCCATATAGCCCCCGCTGTTCGACTTACAAACACAGGCACAGTACTGACAAACCCATACACCTCCTCTGAAATACCCATAGTTGCTAGGGCTGTCTCCGAACTCATTACACCCTCCAAAGTCAGAGCTGTAATTTCGCCATCAAGGGCAGCGAGGGCTTCTCCAGATAAAATAGCTTCTGCCGAGAGTCCCGTAAGGGTAGACACTTCAGCTAATCCCTCGATGAGGTCTACTAGAATAGTCAGTGCGGCTCCCATTTTGAAAATTCACTTACTTGATCAGCTTCAGAAGATGGCGGAGGGCCTCCAACACAGTAATTTTCCTCCCGACTCTTAAAATAGAAAATGTCAAGTCAGTTAAGCAGGAAGTGGACTAACTGACGCAGCTGGCCGTGCGACATCCTCTTTTAATTAGTTGCTAGGCAACGCCCTCCAGAGGGCGTGTGGTTTTGCAAGAGGAAGCAAAAGCCTCTCCACCCAGGCCTAGAATGTTTCCACCCAATCATTACTATGACAACAGCTGTTTTTTTTAGTATTAAGCAGAGGCCGGGGACCCCTGGGCCCGCTTACTCTGGAGAAAAAGAAGAGAGGCATTGTAGAGGCTTCCAGAGGCAACTTGTCAAAACAGGACTGCTTCTATTTCTGTCACACTGTCTGGCCCTGTCACAAGGTCCAGCACCTCCATACCCCCTTTAATAAGCAGTTTGGGAACGGGTGCGGGTCTTACTCCGCCCATCCCGCCCCTAACTCCGCCCAGTTCCGCCCATTCTCCGCCCCATGGCTGACTAATTTTTTTTATTTATGCAGAGGCCGAGGCCGCCTCGGCCTCTGAGCTATTCCAGAAGTAGTGAGGAGGCTTTTTTGGAGGCCTAGGCTTTTGCAAAAAGCTAATTC

**Signal sequence_HA_FT_A1AT_WT**

Annotation Key :

HA tag

FLAG tag

Human A1AT WT

CMV promoter

CMV enhancer

AmpR

TTGAGATCCTTTTTTTCTGCGCGTAATCTGCTGCTTGCAAACAAAAAAACCACCGCTACCAGCGGTGGTTTGTTTGCCGGATCAAGAGCTACCAACTCTTTTTCCGAAGGTAACTGGCTTCAGCAGAGCGCAGATACCAAATACTGTTCTTCTAGTGTAGCCGTAGTTAGGCCACCACTTCAAGAACTCTGTAGCACCGCCTACATACCTCGCTCTGCTAATCCTGTTACCAGTGGCTGCTGCCAGTGGCGATAAGTCGTGTCTTACCGGGTTGGACTCAAGACGATAGTTACCGGATAAGGCGCAGCGGTCGGGCTGAACGGGGGGTTCGTGCACACAGCCCAGCTTGGAGCGAACGACCTACACCGAACTGAGATACCTACAGCGTGAGCTATGAGAAAGCGCCACGCTTCCCGAAGGGAGAAAGGCGGACAGGTATCCGGTAAGCGGCAGGGTCGGAACAGGAGAGCGCACGAGGGAGCTTCCAGGGGGAAACGCCTGGTATCTTTATAGTCCTGTCGGGTTTCGCCACCTCTGACTTGAGCGTCGATTTTTGTGATGCTCGTCAGGGGGGCGGAGCCTATGGAAAAACGCCAGCAACGCGGCCTTTTTACGGTTCCTGGCCTTTTGCTGGCCTTTTGCTCACATGTTCTTTCCTGCGTTATCCCCTGATTCTGTGGATAACCGTATTACCGCCTTTGAGTGAGCTGATACCGCTCGCCGCAGCCGAACGACCGAGCGCAGCGAGTCAGTGAGCGAGGAAGCGGAAGAGCGCCCAATACGCAAACCGCCTCTCCCCGCGCGTTGGCCGATTCATTAATGCAGCTGGCACGACAGGTTTCCCGACTGGAAAGCGGGCAGTGAGCGCAACGCAATTAATGTGAGTTAGCTCACTCATTAGGCACCCCAGGCTTTACACTTTATGCTTCCGGCTCGTATGTTGTGTGGAATTGTGAGCGGATAACAATTTCACACAGGAAACAGCTATGACATGATTACGAATTTAATACGACTCACTATAGGGAATTAACTTTCTGGTTTTTCAGTTCCTCGAGGAGCTTTTTGCAAAAGCCTAGGCCTCCAAAAAAGCCTCCTCACTACTTCTGGAATAGCTCAGAGGCCGAGGCGGCCTCGGCCTCTGCATAAATAAAAAAAATTAGTCAGCCATGGGGCGGAGAATGGGCGGAACTGGGCGGAGTTAGGGGCGGGATGGGCGGAGTTAGGGGCGGGACTATGGTTGCTGACTAATTGAGATGCATGCTTTGCATACTTCTGCCTGCTGGGGAGCCTGGGGACTTTCCACACCTGGTTGCTGACTAATTGAGATGCATGCTTTGCATACTTCTGCCTGCTGGGGAGCCTGGGGACTTTCCACACCCTAACTGACACACATTCCACAGCCAAGCTGGCCGCGTACCCAATTCAACAGGCATCTACTGAGTGGACCCAACGCATGAGAGGACAGTGCCAAGCAAGCAACTCAAATGTCCCACCGGTTGGGCATGGCCAGGTAGCCTATGCTGTGTCTGGACGTCCTCCTGCTGGTATAGTTATTTTAAAATCAGAAGGACAGGGAAGGGAGCAGTGGTTCACGCCTGTAATCCCAGCAATTTGGGAGGCCAAGGTGGGTAGATCACCTGAGATTAGGAGTTGGAGACCAGCCTGGCCAATATGGTGAAACCCCGTCTCTACCAAAAAAACAAAAATTAGCTGAGCCTGGTCATGCATGCCTGGAATCCCAACAACTCGGGAGGCTGAGGCAGGAGAATCGCTTGAACCCAGGAGGCGGAGATTGCAGTGAGCCAAGATTGTGCCACTGCACTCCAGCTTGGTTCCCAATAGACCCCGCAGGCCCTACAGGTTGTCTTCCCAACTTGCCCCTTGCTCCATACCACCCCCCTCCACCCCATAATATTATAGAAGGACACCTAGTCAGACAAAATGATGCAACTTAATTTTATTAGGACAAGGCTGGTGGGCACTGGAGTGGCAACTTCCAGGGCCAGGAGAGGCACTGGGGAGGGGTCACAGGGATGCCACCCGGGATCCTCTAGAGTCGACTGGTACCGATATCAGATCTATCGATGAATTCGAGGTTGAGGAGCGAGAGGCAGTTATTACGTACGCAGGTCCTCCTCTGAGATCAGCTTCTGCTCAGAACCTCCAGAACCTCCAGCGTAATCTGGTACGTCGTATGGGTAAGAACCTCCAGAACCTCCCTTGTCATCGTCGTCCTTGTAATCTTTTTGGGTGGGATTCACCACTTTTCCCATGAAGAGGGGAGACTTGGTATTTTGTTCAATCATTAAGAAGACAAAGGGTTTGTTGAACTTGACCTCGGGGGGGATAGACATGGGTATGGCCTCTAAAAACATGGCCCCAGCAGCTTCAGTCCCTTTCTCGTCGATGGTCAGCACAGCCTTATGCACGGCCTTGGAGAGCTTCAGGGGTGCCTCCTCTGTGACCCCGGAGAGGTCAGCCCCATTGCTGAAGACCTTAGTGATGCCCAGTTGACCCAGGACGCTCTTCAGATCATAGGTTCCAGTAATGGACAGTTTGGGTAAATGTAAGCTGGCAGACCTTCTGTCTTCATTTTCCAGGAACTTGGTGATGATATCGTGGGTGAGTTCATTTTCCAGGTGCTGTAGTTTCCCCTCATCAGGCAGGAAGAAGATGGCGGTGGCATTGCCCAGGTATTTCATCAGCAGCACCCAGCTGGACAGCTTCTTACAGTGCTGGATGTTAAACATGCCTAAACGCTTCATCATAGGCACCTTCACGGTGGTCACCTGGTCCACGTGGAAGTCCTCTTCCTCGGTGTCCTTGACTTCAAAGGGTCTCTCCCATTTGCCTTTAAAGAAGATGTAATTCACCAGAGCAAAAACTGTGTCTCTGTCAAGCTCCTTGACCAAATCCACAATTTTCCCTTGAGTACCCTTCTCCACGTAATCGTTGATCTGTTTCTTGGCCTCTTCGGTGTCCCCGAAGTTGACAGTGAAGGCTTCTGAGTGGTACAACTTTTTAACATCCTCCAAAAACTTATCCACTAGCTTCAGGCCCTCGCTGAGGAACAGGCCATTGCCGGTGGTCAGCTGGAGCTGGCTGTCTGGCTGGTTGAGGGTACGGAGGAGTTCCTGGAAGCCTTCATGGATCTGAGCCTCCGGAATCTCCGTGAGGTTGAAATTCAGGCCCTCCAGGATTTCATCGTGAGTGTCAGCCTTGGTCCCCAGGGAGAGCATTGCAAAGGCTGTAGCGATGCTCACTGGGGAGAAGAAGATATTGGTGCTGTTGGACTGGTGTGCCAGCTGGCGGTATAGGCTGAAGGCGAACTCAGCCAGGTTGGGGGTGATCTTGTTGAAGGTTGGGTGATCCTGATCATGGTGGGATGTATCTGTCTTCTGGGCAGCATCTCCCTGGGGATCCTCAGCCAGGGAGACAGTCTTGTCGTCATCGTCTTTGTAGTCAGCCAGGGAGACAGGGACCAGGCAGCACAGGCCTGCCAGCAGGAGGATGCCCCACGAGACAGAAGACGGCATTGTCGATTCACTGTCCCAGGTCAGGACGGTTCACTAAACGAGCTCTGCTTATATAGACCTCCCACCGTACACGCCTACCGCCCATTTGCGTCAACGGGGCGGGGTTATTACGACATTTTGGAAAGTCCCGTTGATTTTGGTGCCAAAACAAACTCCCATTGACGTCAATGGGGTGGAGACTTGGAAATCCCCGTGAGTCAAACCGCTATCCACGCCCATTGGTGTACTGCCAAAACCGCATCACCATGGTAATAGCGATGACTAATACGTAGATGTACTGCCAAGTAGGAAAGTCCCGTAAGGTCATGTACTGGGCATAATGCCAGGCGGGCCATTTACCGTCATTGACGTCAATAGGGGGCGGACTTGGCATATGATACACTTGATGTACTGCCAAGTGGGCAGTTTACCGTAAATACTCCACCCATTGACGTCAATGGAAAGTCCCTATTGGCGTTACTATGGGAACATACGTCATTATTGACGTCAATGGGCGGGGGTCGTTGGGCGGTCAGCCAGGCGGGCCATTTACCGTAAGTTATGTAACGCGGAACTCCATATATGGGCTATGAACTAATGACCCCGTAATTGATTACTATTAATAACTAGTCAATAATCAATGTCAACATGGCGGTCATATTGGACATGAGCCAATATAAATGTACATATTATGATATAGATACAACGTATGCAATGGCCAATAGCCAATATTGATTTATGCTATATAACCAATGACTAATATGGCTAATTGCCAATATTGATTCAATGTATAGATCAGCTTGGCACTGGCCGTCGTTTTACAACGTCGTGACTGGGAAAACCCTGGCGTTACCCAACTTAATCGCCTTGCAGCACATCCCCCTTTCGCCAGCTGGCGTAATAGCGAAGAGGCCCGCACCGATCGCCCTTCCCAACAGTTGCGCAGCCTGAATGGCGAATGGAAATTGTAAGCGTTAATATTTTGTTAAAATTCGCGTTAAATTTTTGTTAAATCAGCTCATTTTTTAACCAATAGGCCGAAATCGGCAAAATCCCTTATAAATCAAAAGAATAGACCGAGATAGGGTTGAGTGTTGTTCCAGTTTGGAACAAGAGTCCACTATTAAAGAACGTGGACTCCAACGTCAAAGGGCGAAAAACCGTCTATCAGGGCGATGGCCCACTACGTGAACCATCACCCTAATCAAGTTTTTTGGGGTCGAGGTGCCGTAAAGCACTAAATCGGAACCCTAAAGGGAGCCCCCGATTTAGAGCTTGACGGGGAAAGCCGGCGAACGTGGCGAGAAAGGAAGGGAAGAAAGCGAAAGGAGCGGGCGCTAGGGCGCTGGCAAGTGTAGCGGTCACGCTGCGCGTAACCACCACACCCGCCGCGCTTAATGCGCCGCTACAGGGCGCGTCAGGTGGCACTTTTCGGGGAAATGTGCGCGGAACCCCTATTTGTTTATTTTTCTAAATACATTCAAATATGTATCCGCTCATGAGACAATAACCCTGATAAATGCTTCAATAATATTGAAAAAGGAAGAGTATGAGTATTCAACATTTCCGTGTCGCCCTTATTCCCTTTTTTGCGGCATTTTGCCTTCCTGTTTTTGCTCACCCAGAAACGCTGGTGAAAGTAAAAGATGCTGAAGATCAGTTGGGTGCACGAGTGGGTTACATCGAACTGGATCTCAACAGCGGTAAGATCCTTGAGAGTTTTCGCCCCGAAGAACGTTTTCCAATGATGAGCACTTTTAAAGTTCTGCTATGTGGCGCGGTATTATCCCGTATTGACGCCGGGCAAGAGCAACTCGGTCGCCGCATACACTATTCTCAGAATGACTTGGTTGAGTACTCACCAGTCACAGAAAAGCATCTTACGGATGGCATGACAGTAAGAGAATTATGCAGTGCTGCCATAACCATGAGTGATAACACTGCGGCCAACTTACTTCTGACAACGATCGGAGGACCGAAGGAGCTAACCGCTTTTTTGCACAACATGGGGGATCATGTAACTCGCCTTGATCGTTGGGAACCGGAGCTGAATGAAGCCATACCAAACGACGAGCGTGACACCACGATGCCTGTAGCAATGGCAACAACGTTGCGCAAACTATTAACTGGCGAACTACTTACTCTAGCTTCCCGGCAACAATTAATAGACTGGATGGAGGCGGATAAAGTTGCAGGACCACTTCTGCGCTCGGCCCTTCCGGCTGGCTGGTTTATTGCTGATAAATCTGGAGCCGGTGAGCGTGGGTCTCGCGGTATCATTGCAGCACTGGGGCCAGATGGTAAGCCCTCCCGTATCGTAGTTATCTACACGACGGGGAGTCAGGCAACTATGGATGAACGAAATAGACAGATCGCTGAGATAGGTGCCTCACTGATTAAGCATTGGTAACTGTCAGACCAAGTTTACTCATATATACTTTAGATTGATTTAAAACTTCATTTTTAATTTAAAAGGATCTAGGTGAAGATCCTTTTTGATAATCTCATGACCAAAATCCCTTAACGTGAGTTTTCGTTCCACTGAGCGTCAGACCCCGTAGAAAAGATCAAAGGATCTTC

**Signal sequence_HA_A1AT_NHK**

Annotation Key :

HA tag

Human A1AT NHK

CMV promoter

CMV enhancer

AmpR

TTGAGATCCTTTTTTTCTGCGCGTAATCTGCTGCTTGCAAACAAAAAAACCACCGCTACCAGCGGTGGTTTGTTTGCCGGATCAAGAGCTACCAACTCTTTTTCCGAAGGTAACTGGCTTCAGCAGAGCGCAGATACCAAATACTGTTCTTCTAGTGTAGCCGTAGTTAGGCCACCACTTCAAGAACTCTGTAGCACCGCCTACATACCTCGCTCTGCTAATCCTGTTACCAGTGGCTGCTGCCAGTGGCGATAAGTCGTGTCTTACCGGGTTGGACTCAAGACGATAGTTACCGGATAAGGCGCAGCGGTCGGGCTGAACGGGGGGTTCGTGCACACAGCCCAGCTTGGAGCGAACGACCTACACCGAACTGAGATACCTACAGCGTGAGCTATGAGAAAGCGCCACGCTTCCCGAAGGGAGAAAGGCGGACAGGTATCCGGTAAGCGGCAGGGTCGGAACAGGAGAGCGCACGAGGGAGCTTCCAGGGGGAAACGCCTGGTATCTTTATAGTCCTGTCGGGTTTCGCCACCTCTGACTTGAGCGTCGATTTTTGTGATGCTCGTCAGGGGGGCGGAGCCTATGGAAAAACGCCAGCAACGCGGCCTTTTTACGGTTCCTGGCCTTTTGCTGGCCTTTTGCTCACATGTTCTTTCCTGCGTTATCCCCTGATTCTGTGGATAACCGTATTACCGCCTTTGAGTGAGCTGATACCGCTCGCCGCAGCCGAACGACCGAGCGCAGCGAGTCAGTGAGCGAGGAAGCGGAAGAGCGCCCAATACGCAAACCGCCTCTCCCCGCGCGTTGGCCGATTCATTAATGCAGCTGGCACGACAGGTTTCCCGACTGGAAAGCGGGCAGTGAGCGCAACGCAATTAATGTGAGTTAGCTCACTCATTAGGCACCCCAGGCTTTACACTTTATGCTTCCGGCTCGTATGTTGTGTGGAATTGTGAGCGGATAACAATTTCACACAGGAAACAGCTATGACCATGATTACGCCAAGCTCTAGCTAGAGGTCGACGGTATACAGACATGATAAGATACATTGATGAGTTTGGACAAACCACAACTAGAATGCAGTGAAAAAAATGCTTTATTTGTGAAATTTGTGATGCTATTGCTTTATTTGTAACCATTATAAGCTGCAATAAACAAGTTGGGGTGGGCGAAGAACTCCAGCATGAGATCCCCGCGCTGGAGGATCATCCAGCCGGCGTCCCGGAAAACGATTCCGAAGCCCAACCTTTCATAGAAGGCGGCGGTGGAATCGAAATCTCGTGATGGCAGGTTGGGCGTCGCTTGGTCGGTCATTTCGAACCCCAGAGTCCCGCTCAGAAGAACTCGTCAAGAAGGCGATAGAAGGCGATGCGCTGCGAATCGGGAGCGGCGATACCGTAAAGCACGAGGAAGCGGTCAGCCCATTCGCCGCCAAGCTCTTCAGCAATATCACGGGTAGCCAACGCTATGTCCTGATAGCGGTCCGCCACACCCAGCCGGCCACAGTCGATGAATCCAGAAAAGCGGCCATTTTCCACCATGATATTCGGCAAGCAGGCATCGCCATGGGTCACGACGAGATCCTCGCCGTCGGGCATGCGCGCCTTGAGCCTGGCGAACAGTTCGGCTGGCGCGAGCCCCTGATGCTCTTCGTCCAGATCATCCTGATCGACAAGACCGGCTTCCATCCGAGTACGTGCTCGCTCGATGCGATGTTTCGCTTGGTGGTCGAATGGGCAGGTAGCCGGATCAAGCGTATGCAGCCGCCGCATTGCATCAGCCATGATGGATACTTTCTCGGCAGGAGCAAGGTGAGATGACAGGAGATCCTGCCCCGGCACTTCGCCCAATAGCAGCCAGTCCCTTCCCGCTTCAGTGACAACGTCGAGCACAGCTGCGCAAGGAACGCCCGTCGTGGCCAGCCACGATAGCCGCGCTGCCTCGTCCTGCAGTTCATTCAGGGCACCGGACAGGTCGGTCTTGACAAAAAGAACCGGGCGCCCCTGCGCTGACAGCCGGAACACGGCGGCATCAGAGCAGCCGATTGTCTGTTGTGCCCAGTCATAGCCGAATAGCCTCTCCACCCAAGCGGCCGGAGAACCTGCGTGCAATCCATCTTGTTCAATCATGCGAAACGATCCTCATCCTGTCTCTTGATCAGATCCGAAAATGGATATACAAGCTCCCGGGAGCTTTTTGCAAAAGCCTAGGCCTCCAAAAAAGCCTCCTCACTACTTCTGGAATAGCTCAGAGGCAGAGGCGGCCTCGGCCTCTGCATAAATAAAAAAAATTAGTCAGCCATGGGGCGGAGAATGGGCGGAACTGGGCGGAGTTAGGGGCGGGATGGGCGGAGTTAGGGGCGGGACTATGGTTGCTGACTAATTGAGATGCATGCTTTGCATACTTCTGCCTGCTGGGGAGCCTGGGGACTTTCCACACCTGGTTGCTGACTAATTGAGATGCATGCTTTGCATACTTCTGCCTGCTGGGGAGCCTGGGGACTTTCCACACCCTAACTGACACACATTCCACAGAATTAATTCGCGTTAAATTTTTGTTAAATCAGCTCATTTTTTAACCAATAGGCCGAAATCGGCAAAATCCCTTATAAATCAAAAGAATAGACCGAGATAGGGTTGAGTGTTGTTCCAGTTTGGAACAAGAGTCCACTATTAAAGAACGTGGACTCCAACGTCAAAGGGCGAAAAACCGTCTATCAGGGCGATGGCCCACTACGTGAACCATCACCCTAATCAAGTTTTTTGGGGTCGAGGTGCCGTAAAGCACTAAATCGGAACCCTAAAGGGAGCCCCCGATTTAGAGCTTGACGGGGAAAGCCGGCGAACGTGGCGAGAAAGGAAGGGAAGAAAGCGAAAGGAGCGGGCGCTAGGGCGCTGGCAAGTGTAGCGGTCACGCTGCGCGTAACCACCACACCCGCCGCGCTTAATGCGCCGCTACAGGGCGCGTGGGGATACCCCCTAGAGCCCCAGCTGGTTCTTTCCGCCTCAGAAGCCATAGAGCCCACCGCATCCCCAGCATGCCTGCTATTGTCTTCCCAATCCTCCCCCTTGCTGTCCTGCCCCACCCCACCCCCCAGAATAGAATGACACCTACTCAGACAATGCGATGCAATTTCCTCATTTTATTAGGAAAGGACAGTGGGAGTGGCACCTTCCAGGGTCAAGGAAGGCACGGGGGAGGGGCAAACAACAGATGGCTGGCAACTAGAAGGCACAGTCGAGGCTGATCAGCGAGCTCTAGCATTTAGGTGACACTATAGAATAGGGCCCTCTAGATCAAAGAGCGTAATCTGGAACATCGTATGGGTAACCGGTTGTACAGAATTCTGCACGGCCTTGGAGAGCTTCAGGGGTGCCTCCTCTGTGACCCCGGAGGTCAGCCCCATTGCTGAAGACCTTAGTGATGCCCAGTTGACCCAGGACGCTCTTCAGATCATAGGTTCCAGTAATGGACAGTTTGGGTAAATGTAAGCTGGCAGACCTTCTGTCTTCATTTTCCAGGAACTTGGTGATGATATCGTGGGTGAGTTCATTTTCCAGGTGCTGTAGTTTCCCCTCATCAGGCAGGAAGAAGATGGCGGTGGCATTGCCCAGGTATTTCATCAGCAGCACCCAGCTGGACAGCTTCTTACAGTGCTGGATGTTAAACATGCCTAAACGCTTCATCATAGGCACCTTCACGGTGGTCACCTGGTCCACGTGGAAGTCCTCTTCCTCGGTGTCCTTGACTTCAAAGGGTCTCTCCCATTTGCCTTTAAAGAAGATGTAATTCACCAGAGCAAAAACTGTGTCTCTGTCAAGCTCCTTGACCAAATCCACAATTTTCCCTTGAGTACCCTTCTCCACGTAATCGTTGATCTGTTTCTTGGCCTCTTCGGTGTCCCCGAAGTTGACAGTGAAGGCTTCTGAGTGGTACAACTTTTTAACATCCTCCAAAAACTTATCCACTAGCTTCAGGCCCTCGCTGAGGAACAAGCCATTGCCGGTGGTCAGCTGGAGCTGGCTGTCTGGCTGGTTGAGGGTACGGAGGAGTTCCTGGAAGCCTTCATGGATCTGAGCCTCCGGAATCTCCGTGAGGTTGAAATTCAGGCCCTCCAGGATTTCATCGTGAGTGTCAGCCTTGGTCCCCAGGGAGAGCATTGCAAAGGCTGTAGCGATGCTCACTGGGGAGAAGAAGATATTGGTGCTGTTGGACTGGTGTGCCAGCTGGCGGTATAGGCTGAAGGCGAACTCAGCCAGGTTGGGGGTGATCTTGTTGAAGGTTGGGTGATCCTGATCATGGTGGGATGTATCTGTCTTCTGGGCAGCATCTCCCTGGGGATCCTCAGCCAGGGAGACAGGGACCAGGCAGCACAGGCCTGCCAGCAGGAGGATGCCCCACGAGACAGAAGACGGCATGGTGGCGGTACCAAGCTTGGGTCTCCCTATAGTGAGTCGTATTAATTTCGATAAGCCAGTAAGCAGTGGGTTCTCTAGTTAGCCAGAGAGCTCTGCTTATATAGACCTCCCACCGTACACGCCTACCGCCCATTTGCGTCAATGGGGCGGAGTTGTTACGACATTTTGGAAAGTCCCGTTGATTTTGGTGCCAAAACAAACTCCCATTGACGTCAATGGGGTGGAGACTTGGAAATCCCCGTGAGTCAAACCGCTATCCACGCCCATTGATGTACTGCCAAAACCGCATCACCATGGTAATAGCGATGACTAATACGTAGATGTACTGCCAAGTAGGAAAGTCCCATAAGGTCATGTACTGGGCATAATGCCAGGCGGGCCATTTACCGTCATTGACGTCAATAGGGGGCGTACTTGGCATATGATACACTTGATGTACTGCCAAGTGGGCAGTTTACCGTAAATACTCCACCCATTGACGTCAATGGAAAGTCCCTATTGGCGTTACTATGGGAACATACGTCATTATTGACGTCAATGGGCGGGGGTCGTTGGGCGGTCAGCCAGGCGGGCCATTTACCGTAAGTTATGTAACGCGGAACTCCATATATGGGCTATGAACTAATGACCCCGTAATTGATTACTATTAATAACTAGTCAATAATCAATGTCAACGCGTATATCTGGCCCGTACATCGCGAAGCAGCGCAAAACGCCTAACCCTAAGCAGATTCTTCATGCAATTGTCGGTCAAGCCTTGCCTTGTTGTAGCTTAAATTTTGCTCGCGCACTACTCAGCGACCTCCAACACACAAGCAGGGAGCAGATACTGGCTTAACTATGCGGCATCAGAGCAGATTGTACTGAGAGTGCACCATAGGGGATCGGGAGATCTCCCGATCCGTCGACGTCAGGTGGCACTTTTCGGGGAAATGTGCGCGGAACCCCTATTTGTTTATTTTTCTAAATACATTCAAATATGTATCCGCTCATGAGACAATAACCCTGATAAATGCTTCAATAATATTGAAAAAGGAAGAGTATGAGTATTCAACATTTCCGTGTCGCCCTTATTCCCTTTTTTGCGGCATTTTGCCTTCCTGTTTTTGCTCACCCAGAAACGCTGGTGAAAGTAAAAGATGCTGAAGATCAGTTGGGTGCACGAGTGGGTTACATCGAACTGGATCTCAACAGCGGTAAGATCCTTGAGAGTTTTCGCCCCGAAGAACGTTTTCCAATGATGAGCACTTTTAAAGTTCTGCTATGTGGCGCGGTATTATCCCGTATTGACGCCGGGCAAGAGCAACTCGGTCGCCGCATACACTATTCTCAGAATGACTTGGTTGAGTACTCACCAGTCACAGAAAAGCATCTTACGGATGGCATGACAGTAAGAGAATTATGCAGTGCTGCCATAACCATGAGTGATAACACTGCGGCCAACTTACTTCTGACAACGATCGGAGGACCGAAGGAGCTAACCGCTTTTTTGCACAACATGGGGGATCATGTAACTCGCCTTGATCGTTGGGAACCGGAGCTGAATGAAGCCATACCAAACGACGAGCGTGACACCACGATGCCTGTAGCAATGGCAACAACGTTGCGCAAACTATTAACTGGCGAACTACTTACTCTAGCTTCCCGGCAACAATTAATAGACTGGATGGAGGCGGATAAAGTTGCAGGACCACTTCTGCGCTCGGCCCTTCCGGCTGGCTGGTTTATTGCTGATAAATCTGGAGCCGGTGAGCGTGGGTCTCGCGGTATCATTGCAGCACTGGGGCCAGATGGTAAGCCCTCCCGTATCGTAGTTATCTACACGACGGGGAGTCAGGCAACTATGGATGAACGAAATAGACAGATCGCTGAGATAGGTGCCTCACTGATTAAGCATTGGTAACTGTCAGACCAAGTTTACTCATATATACTTTAGATTGATTTAAAACTTCATTTTTAATTTAAAAGGATCTAGGTGAAGATCCTTTTTGATAATCTCATGACCAAAATCCCTTAACGTGAGTTTTCGTTCCACTGAGCGTCAGACCCCGTAGAAAAGATCAAAGGATCTTC

**Preprolactin plasmid**

Annotation Key:

FLAG tag

Preprolactin

CMV promoter

NeoR/KanR

AACAAAATATTAACGCTTACAATTTCCATTCGCCATTCAGGCTGCGCAACTGTTGGGAAGGGCGATCGGTGCGGGCCTCTTCGCTATTACGCCAGCTGGCGAAAGGGGGATGTGCTGCAAGGCGATTAAGTTGGGTAACGCCAGGGTTTTCCCAGTCACGACGTTGTAAAACGACGGCCAGTGCCAAGCTGATCTATACATTGAATCAATATTGGCAATTAGCCATATTAGTCATTGGTTATATAGCATAAATCAATATTGGCTATTGGCCATTGCATACGTTGTATCTATATCATAATATGTACATTTATATTGGCTCATGTCCAATATGACCGCCATGTTGACATTGATTATTGACTAGTTATTAATAGTAATCAATTACGGGGTCATTAGTTCATAGCCCATATATGGAGTTCCGCGTTACATAACTTACGGTAAATGGCCCGCCTGGCTGACCGCCCAACGACCCCCGCCCATTGACGTCAATAATGACGTATGTTCCCATAGTAACGCCAATAGGGACTTTCCATTGACGTCAATGGGTGGAGTATTTACGGTAAACTGCCCACTTGGCAGTACATCAAGTGTATCATATGCCAAGTCCGCCCCCTATTGACGTCAATGACGGTAAATGGCCCGCCTGGCATTATGCCCAGTACATGACCTTACGGGACTTTCCTACTTGGCAGTACATCTACGTATTAGTCATCGCTATTACCATGGTGATGCGGTTTTGGCAGTACACCAATGGGCGTGGATAGCGGTTTGACTCACGGGGATTTCCAAGTCTCCACCCCATTGACGTCAATGGGAGTTTGTTTTGGCACCAAAATCAACGGGACTTTCCAAAATGTCGTAATAACCCCGCCCCGTTGACGCAAATGGGCGGTAGGCGTGTACGGTGGGAGGTCTATATAAGCAGAGCTCGTTTAGTGAACCGTCAGAATTTTGTAATACGACTCACTATAGGGCGGCCGGGAATTCGTCGACTGGATCCGGTACCGAGGAGATCTGATGAACATCAAAGGATCGCCATGGAAAGGGTCCCTCCTGCTGCTGCTGGTGTCAAACCTGCTCCTGTGCCAGAGCGTGGCCCCCTTGCCCATCTGTCCCGGCGGGGCTGCCCGATGCCAGGTGACCCTTCGAGACCTGTTTGACCGCGCCGTCGTCCTGTCCCACTACATCCATAACCTCTCCTCAGAAATGTTCAGCGAATTCGATAAACGGTATACCCATGGCCGGGGGTTCATTACCAAGGCCATCAACAGCTGCCACACTTCTTCCCTTGCCACCCCCGAAGACAAGGAGCAAGCCCAACAGATGAATCAAAAAGACTTTCTGAGCCTGATAGTCAGCATATTGCGATCCTGGAATGAGCCTCTGTATCATCTGGTCACGGAAGTACGTGGTATGCAAGAAGCCCCGGAGGCTATCCTATCCAAAGCTGTAGAGATTGAGGAGCAAACCAAACGGCTTCTAGAGGGCATGGAGCTGATAGTCAGCCAGGTTCATCCTGAAACCAAAGAAAATGAGATCTACCCTGTCTGGTCGGGACTTCCATCCCTGCAGATGGCTGATGAAGAGTCTCGCCTTTCTGCTTATTATAACCTGCTCCACTGCCTACGCAGGGATTCACATAAAATCGACAATTATCTCAAGCTCCTGAAGTGCCGAATCATCCACAACAACAACTGCTAAGATTACAAGGATGACGACGATAAGGTTTAAACGGCCGGCCGCGGTCATAGCTGTTTCCTGAACAGATCCCGGGTGGCATCCCTGTGACCCCTCCCCAGTGCCTCTCCTGGCCCTGGAAGTTGCCACTCCAGTGCCCACCAGCCTTGTCCTAATAAAATTAAGTTGCATCATTTTGTCTGACTAGGTGTCCTTCTATAATATTATGGGGTGGAGGGGGGTGGTATGGAGCAAGGGGCAAGTTGGGAAGACAACCTGTAGGGCCTGCGGGGTCTATTGGGAACCAAGCTGGAGTGCAGTGGCACAATCTTGGCTCACTGCAATCTCCGCCTCCTGGGTTCAAGCGATTCTCCTGCCTCAGCCTCCCGAGTTGTTGGGATTCCAGGCATGCATGACCAGGCTCAGCTAATTTTTGTTTTTTTGGTAGAGACGGGGTTTCACCATATTGGCCAGGCTGGTCTCCAACTCCTAATCTCAGGTGATCTACCCACCTTGGCCTCCCAAATTGCTGGGATTACAGGCGTGAACCACTGCTCCCTTCCCTGTCCTTCTGATTTTAAAATAACTATACCAGCAGGAGGACGTCCAGACACAGCATAGGCTACCTGGCCATGCCCAACCGGTGGGACATTTGAGTTGCTTGCTTGGCACTGTCCTCTCATGCGTTGGGTCCACTCAGTAGATGCCTGTTGAATTGGGTACGCGGCCAGCGGCGAGCGGTATCAGCTCACTCAAAGGCGGTAATACGGTTATCCACAGAATCAGGGGATAACGCAGGAAAGAACATGTGAGCAAAAGGCCAGCAAAAGGCCAGGAACCGTAAAAAGGCCGCGTTGCTGGCGTTTTTCCATAGGCTCCGCCCCCCTGACGAGCATCACAAAAATCGACGCTCAAGTCAGAGGTGGCGAAACCCGACAGGACTATAAAGATACCAGGCGTTTCCCCCTGGAAGCTCCCTCGTGCGCTCTCCTGTTCCGACCCTGCCGCTTACCGGATACCTGTCCGCCTTTCTCCCTTCGGGAAGCGTGGCGCTTTCTCATAGCTCACGCTGTAGGTATCTCAGTTCGGTGTAGGTCGTTCGCTCCAAGCTGGGCTGTGTGCACGAACCCCCCGTTCAGCCCGACCGCTGCGCCTTATCCGGTAACTATCGTCTTGAGTCCAACCCGGTAAGACACGACTTATCGCCACTGGCAGCAGCCACTGGTAACAGGATTAGCAGAGCGAGGTATGTAGGCGGTGCTACAGAGTTCTTGAAGTGGTGGCCTAACTACGGCTACACTAGAAGAACAGTATTTGGTATCTGCGCTCTGCTGAAGCCAGTTACCTTCGGAAAAAGAGTTGGTAGCTCTTGATCCGGCAAACAAACCACCGCTGGTAGCGGTGGTTTTTTTGTTTGCAAGCAGCAGATTACGCGCAGAAAAAAAGGATCTCAAGAAGATCCTTTGATCTTTTCTACGGGGTCTGACGCTCAGTGGAACGAAAACTCACGTTAAGGGATTTTGGTCATGAGATTATCAAAAAGGATCTTCACCTAGATCCTTTTAAATTAAAAATGAAGTTTTAAATCAATCTAAAGTATATATGAGTAACCTGAGGCTATGGCAGGGCCTGCCGCCCCGACGTTGGCTGCGAGCCCTGGGCCTTCACCCGAACTTGGGGGGTGGGGTGGGGAAAAGGAAGAAACGCGGGCGTATTGGCCCCAATGGGGTCTCGGTGGGGTATCGACAGAGTGCCAGCCCTGGGACCGAACCCCGCGTTTATGAACAAACGACCCAACACCGTGCGTTTTATTCTGTCTTTTTATTGCCGTCATAGCGCGGGTTCCTTCCGGTATTGTCTCCTTCCGTGTTTCAGTTAGCCTCCCCCTAGGGTGGGCGAAGAACTCCAGCATGAGATCCCCGCGCTGGAGGATCATCCAGCCGGCGTCCCGGAAAACGATTCCGAAGCCCAACCTTTCATAGAAGGCGGCGGTGGAATCGAAATCTCGTGATGGCAGGTTGGGCGTCGCTTGGTCGGTCATTTCGAACCCCAGAGTCCCGCTCAGAAGAACTCGTCAAGAAGGCGATAGAAGGCGATGCGCTGCGAATCGGGAGCGGCGATACCGTAAAGCACGAGGAAGCGGTCAGCCCATTCGCCGCCAAGCTCTTCAGCAATATCACGGGTAGCCAACGCTATGTCCTGATAGCGATCCGCCACACCCAGCCGGCCACAGTCGATGAATCCAGAAAAGCGGCCATTTTCCACCATGATATTCGGCAAGCAGGCATCGCCATGGGTCACGACGAGATCCTCGCCGTCGGGCATGCTCGCCTTGAGCCTGGCGAACAGTTCGGCTGGCGCGAGCCCCTGATGCTCTTCGTCCAGATCATCCTGATCGACAAGACCGGCTTCCATCCGAGTACGTGCTCGCTCGATGCGATGTTTCGCTTGGTGGTCGAATGGGCAGGTAGCCGGATCAAGCGTATGCAGCCGCCGCATTGCATCAGCCATGATGGATACTTTCTCGGCAGGAGCAAGGTGAGATGACAGGAGATCCTGCCCCGGCACTTCGCCCAATAGCAGCCAGTCCCTTCCCGCTTCAGTGACAACGTCGAGCACAGCTGCGCAAGGAACGCCCGTCGTGGCCAGCCACGATAGCCGCGCTGCCTCGTCTTGCAGTTCATTCAGGGCACCGGACAGGTCGGTCTTGACAAAAAGAACCGGGCGCCCCTGCGCTGACAGCCGGAACACGGCGGCATCAGAGCAGCCGATTGTCTGTTGTGCCCAGTCATAGCCGAATAGCCTCTCCACCCAAGCGGCCGGAGAACCTGCGTGCAATCCATCTTGTTCAATCATGCGAAACGATCCTCATCCTGTCTCTTGATCGATCTTTGCAAAAGCCTAGGCCTCCAAAAAAGCCTCCTCACTACTTCTGGAATAGCTCAGAGGCCGAGGCGGCCTCGGCCTCTGCATAAATAAAAAAAATTAGTCAGCCATGGGGCGGAGAATGGGCGGAACTGGGCGGAGTTAGGGGCGGGATGGGCGGAGTTAGGGGCGGGACTATGGTTGCTGACTAATTGAGATGCATGCTTTGCATACTTCTGCCTGCTGGGGAGCCTGGGGACTTTCCACACCTGGTTGCTGACTAATTGAGATGCATGCTTTGCATACTTCTGCCTGCTGGGGAGCCTGGGGACTTTCCACACCCTAACTGACACACATTCCACAGCTGGTTCTTTCCGCCTCAGGACTCTTCCTTTTTCAATATTATTGAAGCATTTATCAGGGTTATTGTCTCATGAGCGGATACATATTTGAATGTATTTAGAAAAATAAACAAATAGGGGTTCCGCGCACATTTCCCCGAAAAGTGCCACCTGACGCGCCCTGTAGCGGCGCATTAAGCGCGGCGGGTGTGGTGGTTACGCGCAGCGTGACCGCTACACTTGCCAGCGCCCTAGCGCCCGCTCCTTTCGCTTTCTTCCCTTCCTTTCTCGCCACGTTCGCCGGCTTTCCCCGTCAAGCTCTAAATCGGGGGCTCCCTTTAGGGTTCCGATTTAGTGCTTTACGGCACCTCGACCCCAAAAAACTTGATTAGGGTGATGGTTCACGTAGTGGGCCATCGCCCTGATAGACGGTTTTTCGCCCTTTGACGTTGGAGTCCACGTTCTTTAATAGTGGACTCTTGTTCCAAACTGGAACAACACTCAACCCTATCTCGGTCTATTCTTTTGATTTATAAGGGATTTTGCCGATTTCGGCCTATTGGTTAAAAAATGAGCTGATTTAACAAAAATTTAACGCGAATTTT
